# Non-Canonical Targets of HIF1a Drive Cell-Type-Specific Dysfunction

**DOI:** 10.1101/2020.04.03.003632

**Authors:** Kevin C. Allan, Lucille R. Hu, Andrew R. Morton, Marissa A. Scavuzzo, Artur S. Gevorgyan, Benjamin L.L. Clayton, Ilya R. Bederman, Stevephen Hung, Cynthia F. Bartels, Mayur Madhavan, Paul J. Tesar

## Abstract

All mammalian cells sense and respond to insufficient oxygen, or hypoxia, through the activity of hypoxia-inducible factors (HIFs), an evolutionarily conserved family of transcriptional regulators that promote oxygen-independent energy metabolism and angiogenesis. While HIF activation is transiently protective for all cells, prolonged HIF activity drives distinct pathological responses in different tissues. How HIF achieves this pleiotropic effect is largely unknown. Here, we demonstrate that non-canonical targets of HIF1a impair the function of oligodendrocyte progenitor cells (OPCs) to generate oligodendrocytes. Beyond the canonical gene targets shared between all cell types, HIF1a also bound to and activated a unique set of targets in OPCs including *Ascl2* and *Dlx3*. Each of these targets, when ectopically expressed, was sufficient to block oligodendrocyte development through suppression of the key oligodendrocyte regulator *Sox10*. Chemical screening revealed that inhibition of MEK/ERK signaling overcame the HIF1a-mediated block in oligodendrocyte generation by restoring *Sox10* expression without impacting canonical HIF1a activity. Collectively this work defines the mechanism by which chronic HIF1a suppresses oligodendrocyte formation. More broadly, we establish that cell-type-specific HIF1a targets, independent of the canonical hypoxia response, perturb cell function and drive disease in chronic hypoxia.

## INTRODUCTION

The ability to sense and respond to fluctuations in oxygen levels is required to maintain homeostasis in every cell in the body (Kaelin and Ratcliffe, 2008; Semenza, 2012). Insufficient concentrations of molecular oxygen rapidly trigger an evolutionary conserved transcriptional response that enables cell survival in low oxygen by promoting anaerobic metabolism for energy production as well as angiogenesis and erythropoiesis to increase access to local oxygen. While this is initially protective, prolonged activation of this response leads to cellular dysfunction and disease in many tissues. For example, the response to chronic hypoxia blocks white matter formation in premature birth (Scafidi et al., 2014; Volpe, 2009; Volpe et al., 2011), promotes inflammation and insulin resistance in obesity (Lee et al., 2014), and impairs hematopoietic stem cell transplantation capacity (Takubo et al., 2010). This cellular dysfunction has largely been attributed to prolonged activation of the canonical response to low oxygen shared across all cell types; however, it is difficult to explain how activation of a conserved set of hypoxia signature genes can lead to such diverse cellular phenotypes. An alternative unexplored possibility is that cell-type-specific differences in chromatin landscape enable access to unique non-canonical targets, which could account for tissue-specific pathologies.

The response to low oxygen is mediated by hypoxia inducible factors (HIFs), a family of transcription factors that are stabilized under hypoxic conditions in all mammalian cells and are primarily thought to upregulate multiple pathways that adapt cells to low oxygen (Cassavaugh and Lounsbury, 2011; Choudhry and Harris, 2018; Kupferschmidt, 2019). HIFs are heterodimeric complexes consisting of an alpha and beta subunit. In the presence of oxygen, alpha subunits are hydroxylated by prolyl-hydroxylases, allowing for recognition and ubiquitination by von Hippel Lindau (VHL) (Ivan et al., 2001; Jaakkola et al., 2001), and rapid degradation by the proteasome. However, in low oxygen conditions the alpha subunits escape hydroxylation, avoid degradation, and translocate to the nucleus to pair with constitutive beta subunits and regulate gene expression (Cassavaugh and Lounsbury, 2011; Choudhry and Harris, 2018; Semenza, 2007). The HIF1a motif is present more than 1 million times in the genome; however, HIF1a binds to only a small fraction of these sites, which suggests that HIF1a binding is heavily regulated (Schodel et al., 2011; Smythies et al., 2019). Still, the determinants of HIF1a binding in each cell type and whether cell-type-specific targets are functional remain unknown.

The central nervous system (CNS) consumes 20% of the body’s oxygen supply and white matter of the CNS is highly susceptible to hypoxic insults as seen in stroke, vascular dementia, respiratory distress syndromes, premature birth, and subsets of cerebral palsy (Hankey, 2017; Salmaso et al., 2014; Shindo et al., 2016; Volpe, 2009). In fact, chronic HIF1a activity is sufficient to block white matter development (Yuen et al., 2014). White matter of the CNS is formed by oligodendrocytes, which wrap neuronal axons in a lipid-rich protective sheath called myelin, allowing for rapid transmission of action potentials and maintenance of axonal integrity (Chang et al., 2016; Emery, 2010; Nave, 2010). Oligodendrocytes arise from oligodendrocyte progenitor cells (OPCs), which are prevalent in the developing and adult CNS, and HIF1a accumulation has been shown to be sufficient to impair oligodendrocyte formation from OPCs (Jablonska et al., 2016; van Tilborg et al., 2018; Yuen et al., 2014). However, the mechanism of the HIF1a-mediated block in oligodendrocyte formation from OPCs remains unclear. In this study, we use OPCs as an archetypal hypoxia-disease relevant cell type to define the mechanism by which chronic HIF1a drives cell dysfunction.

## RESULTS

### Knockout of VHL models chronic HIF1a accumulation in iPSC-derived OPCs

Defining the mechanisms by which HIF activity perturbs cell function is notoriously challenging as HIFs are rapidly degraded, in minutes, when cells are restored to normoxia. Because of this instability and the low abundance of HIFs, biochemical studies often require extraordinary numbers of cells, which is challenging for hypoxia disease relevant cell types outside of cancer cell lines. To explore the mechanisms underlying the HIF-mediated block in oligodendrocyte development from OPCs, we generated a cellular model of chronic HIF1a accumulation in mouse pluripotent stem cell-derived OPCs, which are uniquely scalable and amenable to genetic manipulation (Hubler et al., 2018; Lager et al., 2018; Najm et al., 2015; Najm et al., 2011). CRISPR-Cas9-mediated knockout of VHL, a central component of the ubiquitin-proteasome system that degrades HIFs (Choudhry and Harris, 2018; Haase, 2009; Rechsteiner et al., 2011), in OPC cultures resulted in stable HIF1a protein accumulation and significant 9-fold and 18-fold activation of downstream hallmark HIF1a targets *Vegfa* and *Bnip3*, respectively (Figures 1A, 1B, S1A and S1B). The response of OPCs to VHL knockout mirrored that of OPCs cultured in hypoxia (1% O_2_), which led to a significant 11-fold increase in *Vegfa* and 26-fold increase in *Bnip3* (Figures 1C and 1D). VHL knockout OPCs were generated with two independent single guide RNAs targeting *Vhl* (sgVhl and sgVhl.2), each of which caused significant decreases in Vhl transcript and protein levels through Cas9-mediated insertion-deletions (in-dels) at the respective target sites compared to control (Cas9 expressing OPCs with no sgRNA) (Figures S1C-S1E). sgVhl OPCs were used for a majority of the data in the study; however, we confirmed key findings in sgVhl.2 OPCs, OPCs treated with physiological hypoxia, primary mouse OPCs exposed to hypoxia, and human OPCs in pluripotent stem cell-derived oligocortical spheroids (Madhavan et al., 2018).

**Figure 1.**
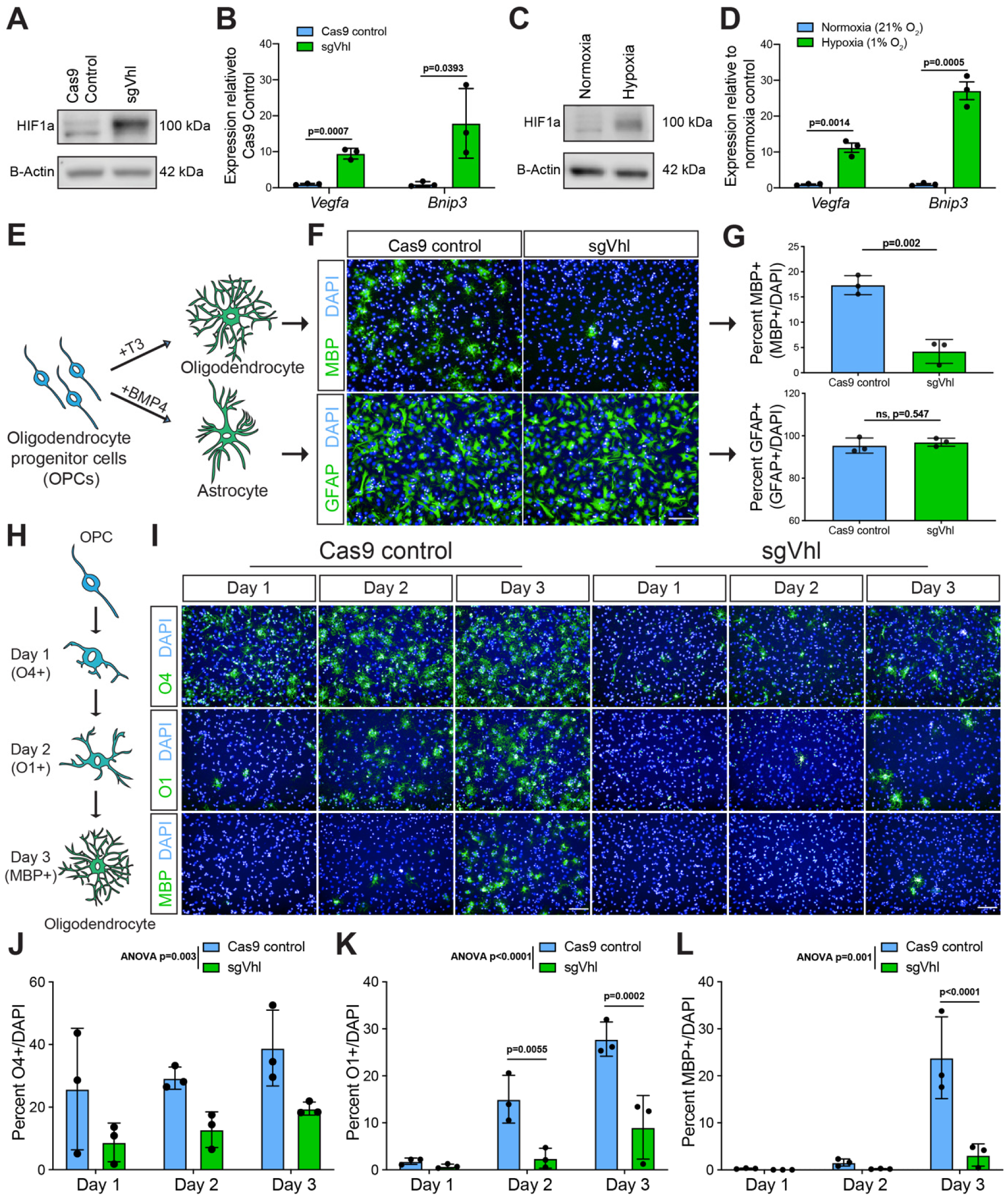
HIF1a Accumulation Impairs the Induction of Oligodendrocytes from OPCs. **(A)** Western blot of HIF1a from nuclear lysates of sgVhl OPCs compared to Cas9 control OPCs with B-Actin as a loading control. Molecular weight is indicated to the right of the blot. **(B)** qRT-PCR of HIF target genes *Vegfa* and *Bnip3* in Cas9 control (in blue) and sgVhl (in green) OPCs normalized to endogenous loading control *Rpl13a*. Data are presented as mean ± SEM from 3 independent biological replicates (experiments) with 4 technical replicates (individual wells) per experiment. p-values were calculated using Student’s two-tailed t-test. **(C)** Western blot of HIF1a from nuclear lysates of OPCs cultured in hypoxia (1% O_2_) compared to normoxia (21% O_2_) with B-Actin as a loading control. Molecular weight is indicated to the right of the blot. **(D)** qRT-PCR of HIF target genes *Vegfa* and *Bnip3* in OPCs cultured in physiological hypoxia (in green) compared to normoxia (in blue) normalized to endogenous loading control *Rpl13a*. Data are presented as mean ± SEM from 3 independent biological replicates (experiments) with 4 technical replicates (individual wells) per experiment. p-values were calculated using Student’s two-tailed t-test. **(E)** Schematic of the two *in vitro* differentiation schemes directing OPCs to either oligodendrocytes through addition of thyroid hormone (T3) or astrocytes through the addition of BMP4. **(F)** Representative images of Cas9 control and sgVhl oligodendrocytes (MBP+ in green) and astrocytes (GFAP+ in green) following 3 days in the respective differentiation media. Nuclei are marked by DAPI (in blue). Scale bar, 100µm. **(G)** Quantification of the percentage of oligodendrocytes (MBP+ cells / DAPI) and astrocytes (GFAP+ cells / DAPI) formed from sgVhl OPCs (in green) compared to Cas9 control OPCs (in blue). Data are presented as mean ± SD from 3 independent biological replicates (experiments) with 6-8 technical replicates (individual wells) per experiment. p-values were calculated using Student’s two-tailed t-test. **(H)** Schematic illustrating acquisition of early (O4), intermediate (O1) and late (MBP) oligodendrocyte markers during the 3-day *in vitro* oligodendrocyte differentiation procedure in response to addition of T3. **(I)** Representative images of early (O4+ in green), intermediate (O1+ in green) and late (MBP+ in green) oligodendrocytes during days 1, 2, and 3 of differentiation with Cas9 control and sgVhl OPCs. Nuclei are marked by DAPI (in blue). Scale bars, 100µm. **(J-L)** Quantification of the percentage of early O4+ (J), intermediate O1+ (K), and late MBP+ (L) oligodendrocytes in sgVhl OPCs (in green) compared to Cas9 control OPCs (in blue) at days 1, 2, and 3 of differentiation. Data are presented as mean ± SD from 3 independent biological replicates (experiments) with 6-8 technical replicates (individual wells) per experiment. To analyze overall differences between Cas9 control and sgVhl OPCs across all timepoints, p-values were calculated using two-way ANOVA (reported as ANOVA p=). To test differences between Cas9 control and sgVhl OPCs at individual timepoints, p-values were calculated using Sidak’s multiple comparisons test. See also Figure S1.

### HIF1a accumulation specifically delays OPC differentiation into oligodendrocytes

HIF1a accumulation in OPCs is sufficient to impair oligodendrocyte formation (Yuen et al., 2014). To test whether HIF1a accumulation is a general or specific inhibitor of OPC differentiation, sgVhl OPCs were stimulated to form either astrocytes (Grinspan et al., 2000), or oligodendrocytes (Baas et al., 1997; Barres et al., 1994; Gao et al., 1998; Najm et al., 2011) (Figure 1E). This revealed a significant 4-fold reduction in oligodendrocyte formation by staining for myelin basic protein (MBP), a marker of mature oligodendrocytes, with no change in astrocyte formation by staining for glial fibrillary acidic protein (GFAP) in sgVhl OPCs compared to control (Figures 1E-1G, and S1F). This suggests HIF1a specifically blocks oligodendrocyte formation from OPCs; however, at what stage HIF1a accumulation impairs the formation of oligodendrocytes is unknown. Staining for early (O4+), intermediate (O1+), and late (MBP+) oligodendrocyte markers throughout the differentiation process demonstrated a significant and delayed acquisition of all oligodendrocyte markers in sgVhl and sgVhl.2 OPCs compared to control (Figures 1H-1L, and S1G-S1I). These data suggest that HIF1a accumulation impairs early OPC differentiation, thereby delaying the formation of oligodendrocytes and ultimately myelin.

### HIF1a binds proximal to promoters and indirectly suppresses *Sox10* expression in OPCs

To delineate the gene targets of HIF1a in OPCs responsible for blocking oligodendrocyte development, chromatin-linked immunoprecipitation sequencing (ChIP-seq) was used to map its genome-wide chromatin binding profile. Utilizing 100 million control and sgVhl OPCs for HIF1a ChIP-seq identified 503 high-stringent peaks (FDR<0.001) in sgVhl OPCs with clear enrichment proximal to the annotated transcription start site (TSS), which agrees with HIF1a as a promoter centric transcription factor (Schodel et al., 2011; Smythies et al., 2019) (Figures 2A and S2A). HIF1a was enriched at canonical target genes *Vegfa* and *Bnip3* and globally peaks were enriched for HIF motifs and motifs of transcription factors that have been shown to interact with HIF1a including Sp1, c-Myc, and Bmal1 (Huang, 2008; Kaluz et al., 2003; Wu et al., 2017) (Figures S2A and S2B). HIF1a was not found proximal to *Sirt1, Wnt7a*, or *Wnt7b* (Figures S2C-S2E), which have previously been suggested as putative HIF effectors in OPCs (Jablonska et al., 2016; Yuen et al., 2014). Moreover, neither sgVhl OPCs nor wild type OPCs exposed to hypoxia exhibited increased expression of *Sirt1, Wnt7a*, or *Wnt7b* transcripts, suggesting that other targets are likely functioning to block oligodendrocyte formation (Figures S2F-S2I) (Zhang et al., 2020).

**Figure 2.**
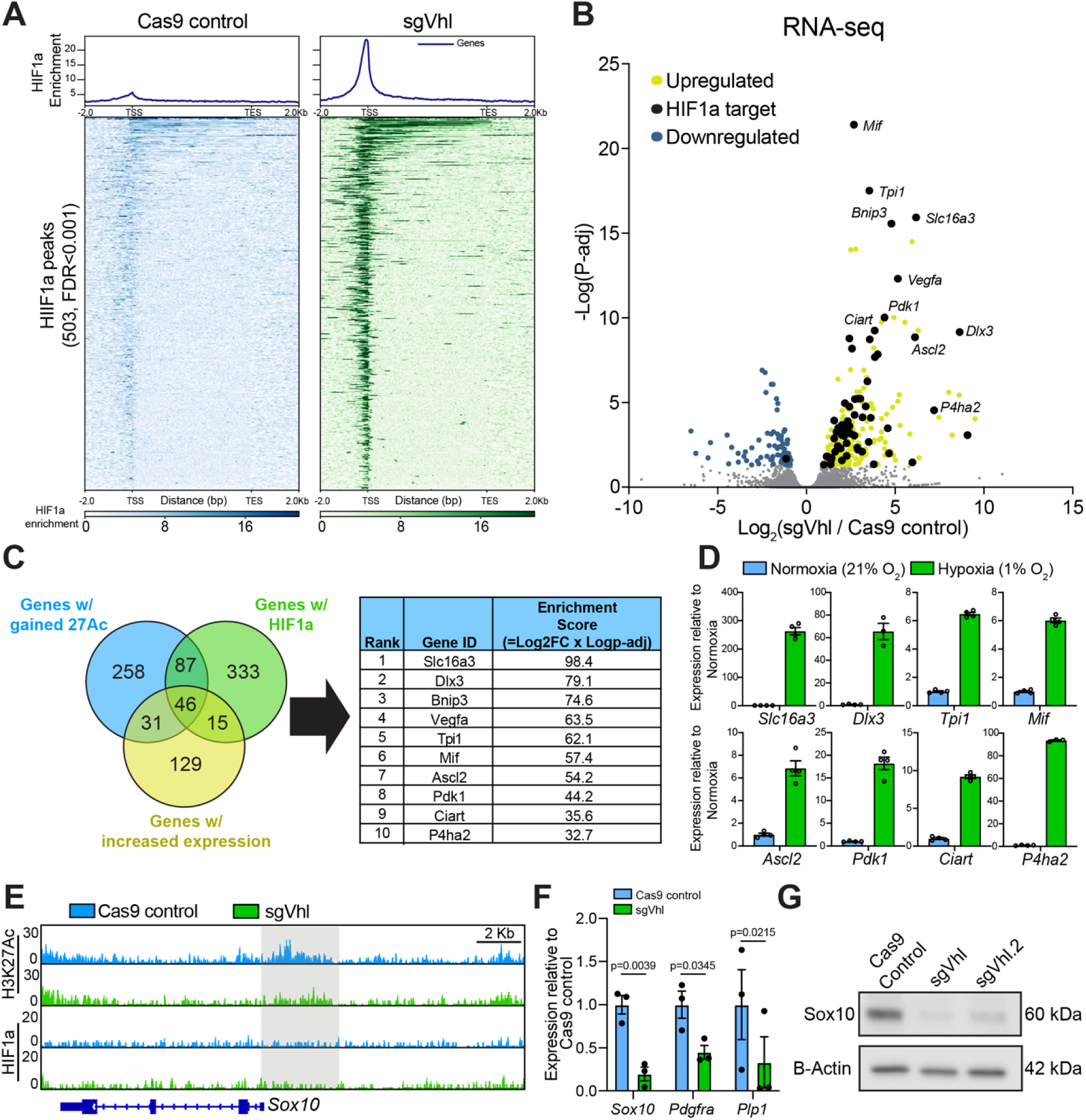
HIF1a Accumulation Suppresses Sox10 Expression. **(A)** Aggregate binding profile and heatmap of the 503 HIF1a peaks called by MACS2 (narrow peaks, FDR<0.001) in sgVhl OPCs within 2Kb of the transcription start site (TSS) and transcription end site (TES) of the closest expressed gene in both Cas9 control and sgVhl OPCs normalized to input. **(B)** Volcano plot of genes that significantly increase (in yellow) and decrease (in blue) with direct targets of HIF1a in black in sgVhl compared to Cas9 control OPCs (p-adj <0.05). Gray dots are genes not significantly different between conditions. Genes indicated in italics represent examples of top direct HIF1a targets in OPCs. Data are from 3 biological replicates (independent samples). **(C)** Venn diagram highlighting top functional HIF1a targets in OPCs by overlapping genes with proximal HIF1a binding (FDR<0.001, in green), significant gains in H3K27ac (FDR<0.1, in blue) and significantly increased expression (P-adj<0.05, in purple). Genes were then ranked by their enrichment score, which is the product of the Log_2_(Fold change in gene expression) and −Log(P-adj). The top 10 targets are displayed in the table ranked by their respective enrichment scores. **(D)** qRT-PCR of 8 top HIF1a target genes (see Figure 1D for *Vegfa* and *Bnip3*) in OPCs treated with hypoxia (1% O_2_, in green) compared to normoxia (21% O_2_, in blue) normalized to endogenous loading control *Rpl13a*. Data are presented as mean ± SEM from 3-4 technical replicates (individual wells) per condition. **(E)** Genome browser view of H3K27ac and HIF1a signals in Cas9 control (in blue) and sgVhl (in green) OPCs normalized to input at the locus for *Sox10* with a reduction in H3K27ac proximal to the *Sox10* promoter highlighted in gray. Scale bar, 2Kb. **(F)** qRT-PCR of *Sox10* and downstream Sox10 target genes *Pdgfra* and *Plp1* in sgVhl OPCs (in green) compared to Cas9 control OPCs (in blue) normalized to endogenous loading control *Rpl13a*. Data are presented as mean ± SEM from 3 biological replicates (independent experiments) with 3-4 technical replicates (individual wells) per experiment. p-values were calculated using Student’s two-tailed t-test. **(G)** Western blot of nuclear lysates for SOX10 in sgVhl and Vhl.2 OPCs compared to Cas9 control OPCs relative to B-Actin loading control. Molecular weight is indicated to the right of the blot. See also Figure S2.

To determine the functional targets of HIF1a, we performed RNA-seq of sgVhl and control OPCs. Overlapping transcripts that significantly changed between sgVhl and control OPCs (P-adj<0.05) with direct targets of HIF1a revealed that HIF1a directly bound to 61 significantly increased genes and only 1 significantly decreased gene (Figure 2B), consistent with the role of HIF1a as a transcriptional activator (Dengler et al., 2014; Guimaraes-Camboa et al., 2015). ChIP-seq for a marker of active chromatin, H3K27Ac (Creyghton et al., 2010), in sgVhl and control OPCs mirrored these findings with a greater number of regions exhibiting a significant gain of H3K27Ac in sgVhl OPCs compared to control (FDR<0.1), and these regions were enriched for HIF motifs (Figures S2J and S2K). To define the top functional targets of HIF1a in OPCs, we overlapped direct HIF1a targets (FDR<0.001) with genes that exhibited both increased transcription (P-adj<0.05) and increased H3K27Ac (FDR<0.1) in sgVhl OPCs compared to control (Figure 2C). Hits were independently validated in OPCs treated with hypoxia, which showed that all of our top 10 HIF1a targets were significantly upregulated compared to normoxia (Figures 1D and 2D).

Of note, HIF1a did not bind proximal to any master regulators of oligodendrocyte development, such as Sox10, Olig2, and Nkx2.2. However, the proximal promoter region of *Sox10*, a basic helix loop helix (bHLH) transcription factor required for formation of oligodendrocytes from OPCs (Stolt et al., 2004; Stolt et al., 2002), showed a reduction of H3K27Ac in sgVhl OPCs (Figure 2E). The decrease in H3K27Ac correlated with a reduction of Sox10 mRNA and protein as well as reduction in expression of downstream Sox10 target genes, *Plp1* and *Pdgfra*, in sgVhl OPCs and primary OPCs treated with hypoxia (1% O_2_) (Figures 2F, 2G, and S2L-S2N). These data suggest that HIF1a activates gene targets that may ultimately impair expression of *Sox10* to block oligodendrocyte development.

### Chromatin accessibility and cell-type-specific transcription factors define non-canonical HIF1a targets

Whether sustained activation of unique cell-type-specific HIF1a target genes drive cellular dysfunction in hypoxia is unknown. To categorize canonical and cell-type-specific targets of HIF1a, we overlapped HIF1a targets in OPCs with HIF1a targets from the limited number of publicly available datasets derived from other mouse cell types including melanocytes (Loftus et al., 2017), T-cells (Ciofani et al., 2012), and embryonic heart (Guimaraes-Camboa et al., 2015). This analysis identified 51 genes that were HIF1a targets across all 4 cell types (“HIF1a core targets”), 152 genes that were HIF1a targets only in OPCs based on these datasets (“OPC-specific HIF1a targets”) and 2250 genes that were specific to either heart, T-cells, or melanocytes (“Other tissue-specific HIF1a targets”) (Figure 3A). Both core and OPC-specific HIF1a target genes collectively increased in expression in sgVhl OPCs compared to control, and both of these gene sets showed a significantly greater increase in expression compared to the other tissue specific HIF1a targets in sgVhl OPCs (Figures 3B and S3A).

**Figure 3.**
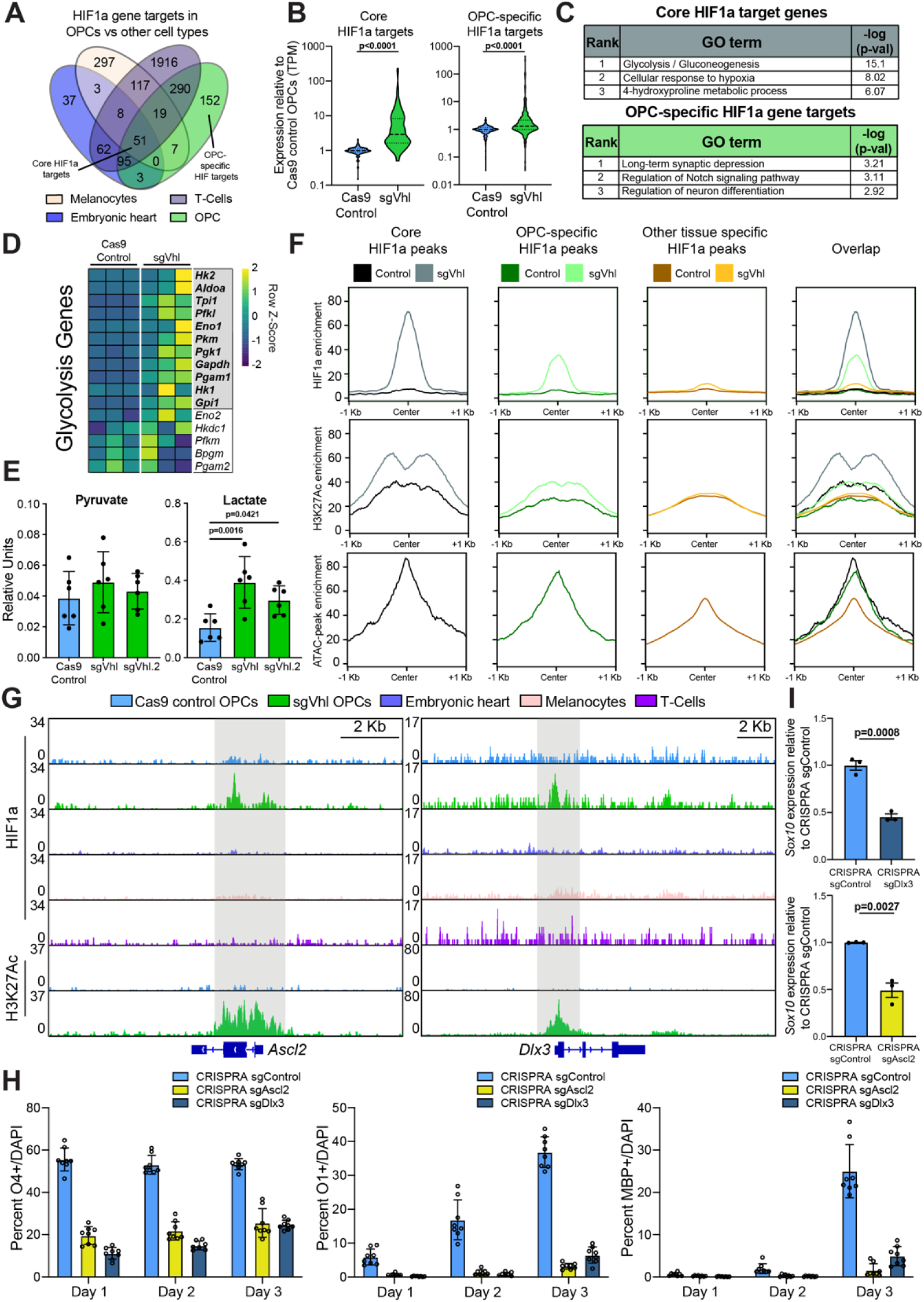
Ectopic Expression of OPC-specific HIF1a Targets Downregulates Sox10 and Impairs Oligodendrocyte Differentiation. **(A)** Venn diagram of direct HIF1a target genes from publicly available HIF1a ChIP-seq datasets in diverse mouse tissues overlapped with HIF1a targets in OPCs (using MACS2, narrowpeaks, FDR<0.001, limited to genes within 5Kb of significant HIF1a peaks). **(B)** Violin plots of compiled expression data (TPM) normalized to Cas9 control OPCs for genes in core and OPC-specific HIF1a target categories in Cas9 control (in blue) and sgVhl (in green). Bold dashed line represents the median with the thin dashed lines representing the upper and lower quartiles. p-values were calculated using Mann Whitney two-tailed t-test. **(C)** Gene ontology (GO) analysis of genes that are core targets of HIF1a in all 4 tissue types as well as genes that are targets of HIF1a specifically in OPCs. Table shows the rank of the GO term along with –log(p-value). **(D)** Heatmap of row normalized expression of genes in the glycolysis pathway between Cas9 control and sgVhl OPCs. Genes with their names bolded and in the gray box are direct targets of HIF1a and exhibit increased expression in OPCs. **(E)** Targeted metabolomics for pyruvate and lactate in both sgVhl and sgVhl.2 OPCs (both in green) compared to Cas9 control OPCs (in blue). Data are presented in relative units as mean ± SD from 6 biological replicates (independent experiments) with one technical replicate (individual well) per experiment. p-values were calculated using one-way ANOVA with Dunnett’s multiple comparisons test. **(F)** Aggregate plots of HIF1a and H3K27ac enrichment in Cas9 control and sgVhl OPCs as well as open chromatin (defined by ATAC-seq) enrichment in normal, non-transduced OPCs at core HIF1a peaks shared by all 4 tissue types, OPC-specific HIF1a peaks, and other tissue-specific HIF1a peaks (combination of cell-type-specific peaks from heart, T-cells, and melanocytes). **(G)** Genome browser view showing HIF1a enrichment at *Ascl2* and *Dlx3* loci in sgVhl OPCs (in green) compared to Cas9 control OPCs (in blue), embryonic heart (in light purple), melanocytes (in beige) and T-cells (in magenta) normalized to input. H3K27ac enrichment is also shown in sgVhl OPCs (in green) compared to Cas9 control OPCs (in blue). HIF1a and H3K27ac accumulation in sgVhl OPCs are highlighted in gray (Scale bars are 2Kb). **(H)** Quantification of the percentage of early O4+, intermediate O1+, and late MBP+ oligodendrocytes in both sgAscl2 (in yellow) and sgDlx3 (in dark blue) CRISPRA OPCs compared to CRISPRA sgControl (in light blue) OPCs at day 1, 2 and 3 of differentiation. Data are presented as mean ± SD of 6-8 technical replicates (individual wells) per condition. **(I)** qRT-PCR of *Sox10* in CRISPRA sgAscl2 (in yellow) and CRISPRA sgDlx3 (in dark blue) compared to CRISPRA sgControl OPCs normalized to endogenous loading control *Rpl13a*. Data are presented as mean ± SEM from 3 biological replicates (independent experiments) with 3-4 technical replicates (individual wells) per experiment. p-values were calculated using Student’s two-tailed t-test. See also Figure S3.

Gene ontology (GO) analysis of core HIF1a target genes showed enrichment for metabolic and hypoxia pathways, which agrees with HIF1a’s canonical role to promote glycolysis in a majority of cell types (Figure 3C) (Choudhry and Harris, 2018; Majmundar et al., 2010; Miska et al., 2019; Nagao et al., 2019). In fact, more than half of the enzymes in the glycolysis pathway were direct HIF1a targets in OPCs, and both sgVhl and sgVhl.2 OPCs exhibited a 2-fold increase in levels of the glycolysis byproduct, lactate, compared to control OPCs (Figures 3D and 3E). Interestingly, GO analysis for cell-type-specific HIF1a targets demonstrated enrichment for pathways separate from the canonical hypoxic response and related to the tissue of origin (Figure S3B). In particular, GO analysis for OPC-specific HIF1a targets showed enrichment for neural development and differentiation pathways (Figure 3C).

To better understand the determinants of the cell type specificity of HIF1a binding profiles, we compared the chromatin landscape in OPCs at core, OPC-specific, and other tissue-specific HIF1a peaks (Figure S3C). The core and OPC-specific HIF1a peaks exhibited a greater enrichment for HIF1a, H3K27Ac and open chromatin (defined by ATAC-seq regions in non-transduced OPCs) compared to other tissue-specific HIF1a sites (Figure 3F). This agrees with previous findings that HIF1a preferentially binds open and active chromatin (Smythies et al., 2019; Xia and Kung, 2009). However, there was a subset of other tissue specific HIF1a peaks enriched for open and active chromatin that lacked HIF1a binding in OPCs, suggesting that chromatin accessibility and activity were not the sole predictors of HIF1a binding. Motif enrichment analysis under cell-type-specific HIF1a peaks demonstrated an enrichment for lineage defining transcription factors such as the Mitf family in melanocytes (Levy et al., 2006), Nkx2 family in embryonic heart (Bartlett et al., 2010), and basic-helix-loop helix (bHLH) motifs in OPCs (Figure S3D). Specifically, the motif for Olig2, a lineage-defining bHLH transcription factor in OPCs (Yu et al., 2013), was highly enriched (in the top 5% of motifs) under OPC-specific HIF1a peaks, whereas the Olig2 motif was not enriched under any other tissue-specific HIF1a peaks (Figure S3D). Collectively, these data suggest that, outside of canonical HIF1a targets, chromatin accessibility and interaction with lineage defining transcription factors determine HIF1a’s unique non-canonical targets in each cell type.

### Cell-type-specific targets of HIF1a suppress oligodendrocyte formation and *Sox10* expression

Out of the top 10 targets of HIF1a in OPCs, *Ascl2* and *Dlx3* were the only OPC-specific HIF1a targets (Figure 2C). Both are transcription factors that regulate differentiation of somatic stem cells in the periphery and are not normally expressed by any cell type in the mouse CNS (Tabula Muris et al., 2018; Zhang et al., 2014); however, both accumulate at the protein level in sgVhl OPCs compared to control (Figures S3E-S3G). This is reflected by the lack of active chromatin at *Ascl2* and *Dlx3* loci in control OPCs; however, both demonstrate robust HIF1a peaks proximal to their promoters specifically in sgVhl OPCs compared to heart, melanocytes and T-cells along with gained H3K27Ac in sgVhl OPCs compared to control OPCs (Figure 3G). Activation of both *Ascl2* and *Dlx3* transcripts also validated in primary OPCs treated with hypoxia (1% O_2_) compared to normoxia (Figure S3H). In fact, ASCL2 was induced *in vivo* in the brains of mouse pups reared in chronic hypoxia (10% O_2_) and this correlated with a decrease in white matter proteins MBP and MAG compared to normoxia reared controls (Figures S3I and S3J).

Ectopic expression of Dlx3 and Ascl2 in OPCs using CRISPR activation (CRISPRA) technology was sufficient to impair the acquisition of early (O4+), intermediate (O1+), and late (MBP+) oligodendrocyte markers across the course of differentiation (Figure 3H). Moreover, ectopic expression of Dlx3 and Ascl2 also led to a significant reduction in *Sox10* expression (Figures 3I and S3K), whereas activation of shared HIF1a core targets, *Slc16a3* and *Vegfa*, did not (Figures S3K and S3L). These data demonstrate that non-canonical HIF1a targets are sufficient to impair oligodendrocyte formation and emphasize that cell-type-specific targets of HIF1a play a role in driving cellular dysfunction.

### Chemical inhibition of MEK/ERK increases oligodendrocyte formation from sgVhl OPCs

To identify potential pathways that could overcome the HIF1a-indcued cellular pathology of OPCs, we tested a library of 1753 bioactive compounds for the ability to increase the formation of MBP+ oligodendrocytes relative to DMSO treated sgVhl OPCs, which exhibited a consistent differentiation deficit compared to DMSO treated control OPCs (Figures 4A and S4A-S4D). Compounds that were non-toxic (fold change in total cell number > 0.7 relative to DMSO treated sgVhl OPCs) and enhanced the number and percentage of MBP+ oligodendrocytes (fold change > 3 relative to DMSO treated sgVhl OPCs) were considered primary hits (Figure S4E). MEK inhibitors were enriched among the primary hits and, as a class, demonstrated a significant increase in oligodendrocyte formation compared to all other non-toxic compounds tested in the primary screen (Figures 4B, 4C, and S4F). Interestingly, drugs previously identified to enhance oligodendrocyte formation, such as miconazole and clemastine (Mei et al., 2014; Najm et al., 2015), were not enriched as hits in this screen, highlighting the ability of this screen to identify context-specific modulators of the differentiation block imposed by HIF1a.

**Figure 4.**
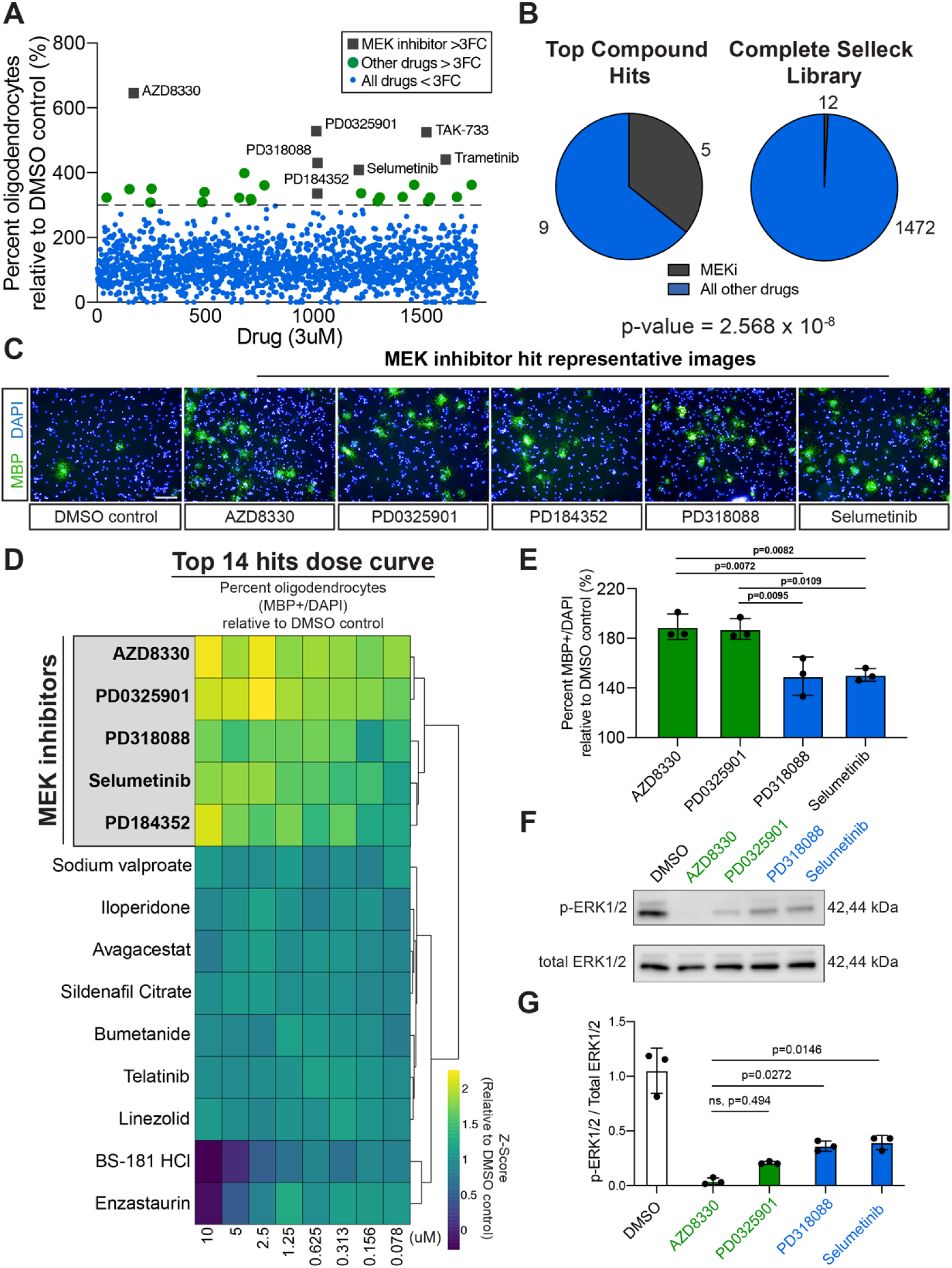
Chemical Inhibition of MEK/ERK Increases Oligodendrocyte Formation from sgVhl OPCs. **(A)** Primary bioactives library screen showing the effect of 1753 molecules on percentage of oligodendrocytes (MBP+ cells/ total DAPI) formed by sgVhl OPCs relative to DMSO treated sgVhl OPCs. Anything above the dotted line represents a greater than 3-fold change increase in percentage of oligodendrocytes from DMSO (green dots). MEK inhibitors are highlighted as gray boxes with their respective drug names. **(B)** Pie charts of the number of non-toxic MEK inhibitors (in dark gray) compared to other classes of drugs (in blue) within top compound hits compared to their prevalence in the non-toxic compounds of the Selleck library. p-value was calculated using hypergeometric analysis. **(C)** Representative ICC images of oligodendrocytes (MBP+ in green) from the primary drug screen of the 5 top MEK inhibitor hits along with the DMSO negative control. Nuclei are marked by DAPI (in blue). Scale bars, 100µm. **(D)** Heatmap representation of the top 14 hits in an 8-point dose curve in 2-fold dilutions from 10µM to 78nM showing the fold change in the percentage (MBP+/total DAPI) of oligodendrocytes relative to DMSO treated sgVhl OPCs. The heatmap is shown as row Z-score, and rows are sorted by unsupervised hierarchical clustering with columns in order from high (10µM) to low dose (78nM). MEK inhibitors are highlighted in gray and bolded. Data are presented as the mean for each drug at each dose from 3 separate dose curve plates. **(E)** Collapsing all tested doses into one overall average shows the ability of each MEK inhibitor to increase the formation of oligodendrocytes (MBP+/DAPI) relative to DMSO treated sgVhl OPCs. AZD8330 and PD0325901 are shown in green representing the most effective drugs, while PD318088 and Selumetinib are shown in blue as slightly less effective drugs. Data are presented as the mean ± SD of the averages of all 8 doses for each drug from 3 separate dose curve plates. p-values were calculated using a one-way ANOVA with Tukey’s multiple comparisons test. **(F)** Representative Western blot for phosphorylated ERK1/2 (p-ERK1/2) relative to total ERK1/2 from whole cell lysates of sgVhl OPCs incubated with 100nM of AZD8330, PD0325901, PD318088 or Selumetinib for 30 minutes. Molecular weight is indicated to the right of the blot. **(G)** Quantification of the ratio of phosphorylated ERK1/2 relative to total ERK1/2 for AZD8330 and PD0325901 (both in green) and PD318088 and Selumetinib (both in blue). Data are presented as mean ± SD from 3 biological replicates (independent experiments) with a single technical replicate per experiment. p-values were calculated using one-way ANOVA with Dunnett’s multiple comparisons test. See also Figure S4.

To identify compounds that were effective across a range of doses, we performed an 8-point dose curve from 10µM to 78nM of the top 14 hits. Performing unbiased hierarchical clustering of the results revealed that all 5 MEK inhibitors clustered together and led to a pronounced increase in oligodendrocyte formation relative to DMSO treated sgVhl OPCs, with AZD8330 and PD0325091 outperforming PD318088 and Selumetinib (Figures 4D, 4E, and S4G). The ability of these different MEK inhibitors to increase oligodendrocyte formation correlated with their on-target IC50 values for MEK1 and MEK2 as well as their on-target ability to reduce ERK1/2 phosphorylation in sgVhl OPCs (Figures 4F, 4G, and S4H). Performance of an 8-point dose curve consisting of 12 drugs that each inhibit a potential downstream target of MEK revealed that ERK1/2 inhibitors led to the greatest increase in oligodendrocyte formation compared to the other classes of drugs tested (Shaul and Seger, 2007; Yohe et al., 2018) (Figures S4I and S4J). Collectively, these results demonstrate that chemical inhibition of MEK/ERK acts as a node of intervention to increase the formation of oligodendrocytes in the context of HIF1a accumulation.

### MEK/ERK inhibition drives *Sox10* expression without changing HIF1a activity

We next asked whether these drugs increased differentiation of sgVhl OPCs by inhibiting HIF1a signaling or circumventing HIF1a by driving genes critical for oligodendrocyte development. Treating OPCs with 300nM AZD8330, our most potent MEK inhibitor (Figures 4F and 4G), for 14 hours led to no change in HIF1a translocation to the nucleus compared to DMSO treated sgVhl OPCs (Figure 5A). Performing RNA-seq on control and sgVhl OPCs treated with either DMSO or 300nM AZD8330 for 14 hours mirrored these results and demonstrated that AZD8330 treatment did not change HIF1a target gene expression, such as *Ascl2* and *Dlx3*, in sgVhl OPCs compared to DMSO (Figures 5B, 5C, and S5A). This demonstrates that MEK inhibitor treatment does not directly counter HIF1a activity, but rather circumvents the effect of HIF1a accumulation by increasing oligodendrocyte differentiation despite persistent HIF signaling.

**Figure 5.**
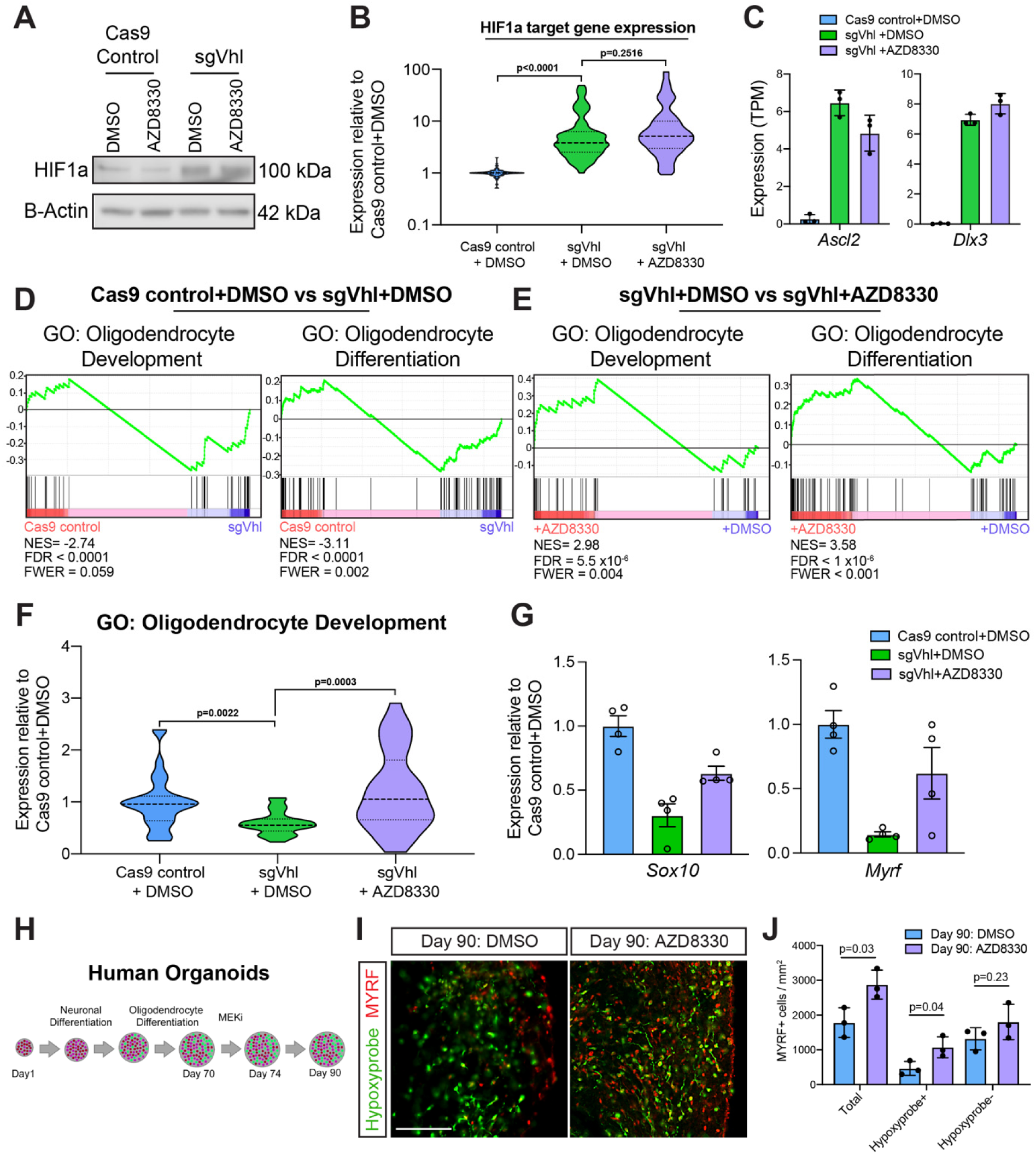
MEK Inhibitors Restore *Sox10* Expression without Changing HIF1a activity in OPCs. **(A)** Western blot of nuclear lysates for HIF1a in sgVhl OPCs with 24hrs of 300nM AZD8330 treatment compared to DMSO with B-Actin as a loading control. Molecular weight is indicated to the right of the blot. **(B)** Violin plot of normalized number of transcripts (TPM values normalized to Cas9 control+DMSO) in Cas9 control+DMSO (in blue), sgVhl + DMSO (in green), and sgVhl + AZD8330 (in purple) relative to Cas9 control + DMSO of genes that were previously shown to be direct targets of HIF and increase in sgVhl OPCs compared to control (see Figure 2B). p-values were calculated using the Kruskal Wallis One-Way ANOVA with Dunn’s multiple comparisons test. **(C)** Quantification of the normalized number of transcripts (TPM) for both *Ascl2* and *Dlx3* in Cas9 control+DMSO (in blue), sgVhl+DMSO OPCs (in green) and sgVhl+AZD8330 OPCs (in purple). OPCs were treated with either DMSO or 300nM AZD8330 for 14 hours. Data represent mean ± SD from 3 biological replicates (independent samples) from RNA-seq. **(D)** Gene set enrichment analysis (GSEA) analysis of gene program changes in sgVhl compared to Cas9 control OPCs demonstrates a significant reduction in GO terms for Oligodendrocyte Development (normalized enrichment score/NES = −2.74, FDR<0.0001, FWER p-val = 0.059) and Oligodendrocyte Differentiation (NES = −3.11, FDR<0.0001, FWER p-val = 0.002). **(E)** GSEA analysis of gene program changes in sgVhl + AZD8330 compared to sgVhl + DMSO OPCs demonstrates a significant enrichment in GO terms for Oligodendrocyte Development (normalized enrichment score/NES = 2.98, FDR= 5.5×10^−6^, FWER p-val = 0.004) and Oligodendrocyte Differentiation (NES = 3.58, FDR<1×10^−6^, FWER p-val <0.001). **(F)** Violin plot showing expression of transcripts (TPM values normalized to Cas9 control + DMSO OPCs) associated with GO term Oligodendrocyte Development (GO:0014003) that decrease (FC<0.75) in sgVhl + DMSO OPCs (in green) relative to Cas9 control + DMSO OPCs (in blue) as well as sgVhl OPCs following treatment with 300nM AZD8330. p-values were calculated using the Kruskal Wallis One-Way ANOVA with Dunn’s multiple comparisons test. **(G)** qRT-PCR of *Sox10* and *Myrf* in Cas9 control+DMSO (in blue), sgVhl+DMSO (in green) and sgVhl+AZD8330 OPCs (in purple) normalized to endogenous control *Rpl13a*. OPCs were treated with either DMSO or 300nM AZD8330 for 14 hours. Data are presented as mean ± SEM from 4 technical replicates (individual wells). **(H)** Schematic of human brain oligocortical spheroids treated at days *in vitro* 70 with either DMSO or 300nM AZD8330 for 4 days. The spheroids were then cultured without drug until day 90 when they were incubated with hypoxyprobe, fixed and then sectioned for immunohistochemistry. **(I)** Representative immunohistochemistry images of DIV 90 oligocortical spheroids that had been treated from DIV 70-74 with either DMSO or 300nM AZD8330 for oligodendrocytes (MYRF+ in red) and hypoxic regions (hypoxyprobe in green). Scale bar, 100µM. **(J)** Quantification of oligodendrocytes (MYRF+ / mm^2^) in the whole oligocortical spheroid (total), hypoxic region of the spheroid (hypoxyprobe+), and normoxic region of the spheroid (hypoxyprobe-) in DIV 90 spheroids that had been treated with either DMSO or 300nM AZD8330 from DIV 70-74. Data represent mean ± SD from 3 biological replicates (individual spheroids). p-values were calculated using Student’s two-tailed t-test. See also Figure S5.

Gene set enrichment analysis (GSEA) revealed that AZD8330 treatment of sgVhl OPCs led to an enrichment for “Oligodendrocyte Differentiation” and “Oligodendrocyte Development” pathways, which were normally depleted in sgVhl OPCs compared to control (Figures 5D and 5E). Further supporting this, AZD8330 treatment of sgVhl OPCs led to a significant increase in the subset of genes within the GO term “Oligodendrocyte Development, GO:0014003” that were normally decreased in sgVhl OPCs compared to control (fold change of sgVhl to control OPCs <0.75), such as *Sox10* and *Myrf* (Figures 5F and 5G). We validated the AZD8330-mediated increase in expression of both of these transcription factors, which are critical for oligodendrocyte differentiation, by qPCR (Figure 5G). To confirm that this was a function of impaired MEK/ERK signaling, treatment of sgVhl OPCs with ERK1/2 inhibitors AZD0364 and SCH772984 led to a similar increase in *Sox10* expression compared to DMSO (Figure S5B). Collectively, these data suggest that the reduction of *Sox10* expression by cell-type-specific HIF1a targets is critical for the HIF-mediated block in oligodendrocyte differentiation, such that restoration of *Sox10* expression without altering canonical HIF function restores oligodendrocyte formation.

### MEK/ERK inhibition drives oligodendrocyte formation in hypoxic regions of human oligocortical spheroids

In order to ascertain if these mechanisms are conserved in the human context, we leveraged a previously established method of generating human myelinating cortical spheroids from pluripotent stem cells (Madhavan et al., 2018). The interior of human brain spheroids is hypoxic (Brawner et al., 2017; Giandomenico and Lancaster, 2017), and we hypothesized that these hypoxic regions would inhibit oligodendrocyte formation, which could be overcome using a MEK inhibitor. To test this, we treated oligocortical spheroids with either DMSO or 300nM AZD8330 for 4 days starting at 70 days *in vitro*, immediately following induction of oligodendrocytes. At day 90, spheroids were treated with hypoxyprobe, a chemical used to visualize hypoxic regions less than 1% O_2_ (Pogue et al., 2001), and harvested for analysis (Figure 5H). Immunohistochemistry for oligodendrocytes (MYRF+ cells) and hypoxic regions (defined by hypoxyprobe + staining) demonstrated a significant 3.8-fold reduction in the number of MYRF+ oligodendrocytes in hypoxic regions compared to normoxic regions within the spheroids (Figures 5I and 5J). MEK/ERK inhibition with AZD8330 treatment led to a significant 2.3-fold increase in the number of oligodendrocytes within hypoxic regions of the spheroid (Figures 5I and 5J). Collectively these results show that oxygen tensions shape oligodendrocyte development and that MEK inhibition circumvents the hypoxia-mediated inhibition of oligodendrocyte formation in 3D models of human brain development.

## DISCUSSION

Cells are equipped to translate external cues from the environment into internal signals that ultimately alter transcriptional programs. Molecular oxygen is crucial to support energy production of the cell, and low oxygen upregulates a rapid and conserved transcriptional response mediated largely by HIF transcription factors in all mammalian cells. HIF1a promotes an adaptive response by upregulating oxygen-independent metabolism and increasing blood vessel formation; however, chronic accumulation of HIF1a negatively impacts the function of almost every organ system (Kullmann et al., 2020; Lee et al., 2019; Menendez-Montes et al., 2016; Takubo et al., 2010).

Here, we profiled the genome-wide functional targets of HIF1a in OPCs and found that HIF1a not only binds to canonical hypoxia response genes that are shared across multiple cell types, but also activates a unique set of non-canonical genes in a cell-type-specific manner. In the context of the brain, these non-canonical HIF1a targets that impair oligodendrocyte formation could have implications in the numerous hypoxia driven pathologies of white matter such as diffuse white matter injury of prematurity (Salmaso et al., 2014; van Tilborg et al., 2018), white matter stroke in adults (Hankey, 2017; Marin and Carmichael, 2018) and multiple sclerosis (Graumann et al., 2003; Zeis et al., 2008). More broadly, we suggest that non-canonical HIF1a targets in diverse cell types impact a variety of cell-type-specific functions, such as oligodendrocyte differentiation, heart morphogenesis and T-cell activation. This concept has previously been overlooked as many studies have focused on canonical targets of HIF1a or those that were discovered in immortalized cell lines, which may behave differently in response to HIF1a accumulation.

The same transcription factor can bind to different gene targets in different cell types through interaction with transcriptional machinery unique to each cell type (Mullen et al., 2011; Trompouki et al., 2011). HIF1a binding has been shown to depend on the openness and activation status of chromatin; however, we and others show that chromatin accessibility is not the sole predictor of HIF1a binding (Schodel et al., 2011; Smythies et al., 2019; Xia and Kung, 2009). Our results highlight that HIF1a binds more strongly at core hypoxia response peaks compared to cell-type-specific peaks, which are enriched for open chromatin as well as motifs for lineage-defining transcription factors. This implies that acute HIF1a binds more readily to protective pathways in the immediate response to low oxygen, while chronic HIF1a interacts with lineage defining transcription factors to upregulate targets that ultimately impair development and function in a cell-type-specific manner. These non-canonical targets could represent either a pathological “off-target” effect of sustained HIF1a accumulation or a normal cell-type-specific response to molecular oxygen levels that is coopted in the context of hypoxia disease.

Overall, this work advances our conceptual understanding of the tissue specific response to chronic HIF accumulation and how oxygen tensions regulate tissue physiology and pathology.

## ACKNOWLEDGEMENTS

This research was supported by grants from the National Institutes of Health F30HD096784 (K.C.A), T32NS077888 (K.C.A.), and T32GM007250 (K.C.A) and the New York Stem Cell Foundation (P.J.T. and M.A.S.) as well as institutional support from CWRU School of Medicine and philanthropic support from the Enrile, Peterson, Fakhouri, Long, Goodman, Geller, and Weidenthal families. Additional support was provided by the Small Molecule Drug Development and Genomics core facilities of the CWRU Comprehensive Cancer Center (P30CA043703), the CWRU Light Microscopy Imaging Center (S10-OD016164), and the University of Chicago Genomics Facility. The authors thank Brian Popko for tissue samples (obtained with funding from NIH grant R01NS034939). We are also grateful to C. Baecher-Allan, J. LaManna, K. Xu, D. Adams, Y. Federov, L. Barbar, E. Prendergast, E. Schwarz, P. Scacheri, D. Neu, E. Cohn, A. Saiakhova, Z. Nevin, M. Elitt, and R. Sallari for technical assistance and/or discussion.

## AUTHOR CONTRIBUTIONS

K.C.A. and P.J.T. conceived and managed the overall study. K.C.A. and L.R.H. performed, quantified and analyzed all *in vitro* experiments using mouse OPCs including qPCR, western blot, immunocytochemistry, and generation of CRISPR knockout and CRISPRA OPCs. K.C.A. performed the small molecule screen and dose curve validations. C.F.B. trained K.C.A. to perform ChIP-seq and K.C.A. performed all ChIP-seq experiments in the paper with analysis of data performed by A.R.M and S.H. A.R.M. and M.A.S. assisted with RNA-seq data analysis. I.R.B. performed mass spectroscopy and quantified the data. B.L.C. generated immunopanned *in vivo* derived OPCs and brain tissue samples. M.M. and A.G. performed oligocortical spheroid experiments. M.A.S. and B.L.C contributed key ideas for experimental design and assembly of figures. K.C.A. assembled all figures and performed statistical analyses. K.C.A. and P.J.T. wrote the manuscript with input from all authors.

## DECLARATION OF INTERESTS

P.J.T. is a co-founder and consultant for Convelo therapeutics, which has licensed unrelated patents from Case Western Reserve University. P.J.T and Case Western Reserve University hold equity in Convelo Therapeutics. All other authors have no competing interests.

## METHODS

### OPC preparation and culture

OPCs were generated from the epiblast stem cells (EpiSCs) as previously described (Najm et al., 2011). These EpiSC derived OPCs were sorted to purity by fluorescence activated cell sorting using conjugated CD140a-APC (eBioscience, 17-1401; 1:80) and NG2-AF488 (Millipore, AB5320A4; 1:100) antibodies. Primary mouse OPCs were derived using two methods. In the first method, cerebral cortices were harvested from postnatal day 2 (P2) C57BL/6J pups and dissociated using a Tumor Dissociation Kit (Miltenyi). Cells were then filtered through a 70 µm filter, washed in DMEM/F12, and plated on poly-ornithine (PO) (Sigma, P3655-50MG) and laminin (Sigma, L2020-1MG) coated plates to be expanded, passaged, and used in experiments. The second method follows the immunopanning protocol from P7 C57BL/6J mice (Barres et al., 1992). Primary *in vivo* derived OPCs using either method were pooled from multiple pups such that they are a combination of male and female cells.

All OPCs were grown on PO and laminin coated flasks in growth media consisting of DMEM/F12 supplemented with N2 Max (R&D Systems), B27 (Thermo Fisher), 20ng/mL bFGF (R&D Systems), and 20ng/mL PDGFA (R&D Systems). Media was changed every 48 hours.

### Mouse OPC differentiation to oligodendrocytes and astrocytes

For oligodendrocyte generation, OPCs were seeded at either 40,000 cells per well (96-well plate, Fisher, 167008) or 15,000 cells per well (384-well PDL-coated cell carrier plates, PerkinElmer, 6057500) on plates coated with PO and laminin. Oligodendrocyte differentiation media consisted of DMEM/F12 supplemented with N2 Max, B27, 100ng/mL noggin (R&D, 3344NG050), 100ng/mL IGF-1 (R&D, 291G1200), 10uM cyclic AMP (Sigma, D0260-100MG), 10ng/mL NT3 (R&D, 267N3025) and 40ng/mL T3 (Thyroid hormone, Sigma, T-6397). Cells were analyzed after 3 days unless otherwise noted.

For OPC differentiation to astrocytes, 30,000 OPCs were plated per well in a 96 well plate (Fisher, 167008) containing astrocyte differentiation media described in (Liddelow et al., 2017). This media consisted of a 1:1 (v/v) mixture of neurobasal media and high glucose DMEM supplemented with sodium pyruvate, glutamax, N2 Max, and N-acetyl-cysteine and with growth factors including 20ng/mL bFGF, 5ng/mL Hb-EGF (R&D, 259-HE-050), 10ng/mL CNTF (R&D, 557-NT-010), and 10ng/mL BMP4 (R&D, 314-BP-050) for 3 days.

### *In vitro* hypoxia experiments

OPCs were plated in OPC growth media and then placed into a 2 shelf C-Chamber from BioSpherix (C-274). Oxygen tension was controlled using the ProOx 110 from BioSpherix such that nitrogen gas would flush out oxygen to maintain the chamber at the desired oxygen level. The subchamber was set at 1% O_2_ and cells were cultured for 48 hours unless otherwise noted. Hypoxia treated cells were then rapidly lysed for RNA or protein to minimize degradation of HIFs upon exposure to room air. The normoxic controls were cultured concurrently in the same cell culture incubator containing the BioSpherix C-chamber.

### Immunocytochemistry

For antigens requiring live staining (O1 and O4), antibodies were diluted in N2B27 base media supplemented with 10% Donkey Serum (v/v) (017-000-121, Jackson ImmunoResearch) and then added to cells for 18 minutes at 37°C. Cells were then fixed in cold 4% PFA (Electron microscopy sciences) for 18 minutes at room temperature, washed with PBS, and permeabilized and blocked in blocking solution containing 0.1% Triton X-100 in PBS supplemented with 10% normal donkey serum (v/v) for 30 minutes at room temperature. Primary antibodies were diluted in blocking solution and incubated with samples overnight at 4°C. Primary antibodies used included anti-OLIG2 (1.2µg/mL, Proteintech, 12999-1-AP), anti-MBP (1:100, Abcam, ab7349), anti-O1 (1:50, CCF Hybridoma Core), anti-O4 (1:100, CCF Hybridoma Core), anti-ASCL2 (1:10, EMD Millipore, MAB4417), and anti-GFAP (1:5,000, Dako, Z033401-2). The next day, cells were rinsed with PBS and incubated in blocking solution for one hour with the appropriate secondary antibody conjugated to an Alexa-Fluor (4ug/mL, Thermo Fisher) along with the nuclear stain DAPI (Sigma, 1ug/mL).

### High content imaging and quantification

Both 96-well and 384-well plates were imaged using the Operetta High Content imaging and analysis system (PerkinElmer). For 96-well and 384-well plates, 8 fields and 5-fields were captured from each well at 20x magnification respectively. Images were analyzed with PerkinElmer Harmony and Columbus software as described previously (Hubler et al., 2018; Najm et al., 2015). In brief, nuclei of live cells were identified using a threshold for area of DAPI staining to exclude pyknotic nuclei or debris. To identify oligodendrocytes, each DAPI positive nucleus was expanded by 50% to determine potential intersection with staining of an oligodendrocyte marker (O4/O1/MBP) in a separate channel. Expanded nuclei that intersected O4/O1/MBP staining were scored as oligodendrocytes. Percentage of oligodendrocytes was then calculated by dividing the number of oligodendrocytes by total number of DAPI positive cells per image.

### Generation of CRISPRKO/ CRISPRA OPCs

Guide sequences were curated from the Brie library (Doench et al., 2016) and cloned into the CRISPRv2 backbone (Addgene 52961) (Sanjana et al., 2014) for generating CRISPR-mediated knockout OPCs. Guide sequences were curated from the CRISPRav2 library (Horlbeck et al., 2016) and cloned into the lenti-SAMv2 backbone (Addgene 75112) (Joung et al., 2017) for CRISPR activation. The activation helper plasmid (Addgene 89308) (Joung et al., 2017) was co-transduced for all CRISPR activation OPCs. Plasmids containing cloning sites for the sgRNA were digested and annealed oligomers containing the sgRNA (indicated in the chart below) were then ligated to their respective backbones. Positive bacterial clones were confirmed to have the correct inserted oligomer using Sanger sequencing. HEK293T cells were then transfected using lenti-X shots following the manufacturer’s protocol (Clonetech). The next day, transfection media was switched to OPC growth media without growth factors for virus collection. After 2 days, the media from transfected HEK293T cells was collected, filtered, and supplemented with OPC growth factors PDGF and FGF. This lentivirus-containing media was added to OPCs at a ratio of 1:2 (v/v) with fresh OPC growth media. After 24 hours of incubation with virus, transduced cells were switched to fresh OPC growth media and allowed to recover for 48 hours. OPCs were then selected for 96 hours in OPC growth media supplemented with a lethal dose of puromycin (500ng/mL, Invitrogen) for CRISPR knockout cells or a lethal dose of blasticidin (10µg/mL, Thermo Fisher, A1113903) and hygromycin (100µg/mL, Thermo Fisher, 10687010) for CRISPR activation cells. OPCs were then allowed to recover for at least 24 hours following removal of selection and frozen down in aliquots for future use. For all experiments, the lentivirally transduced CRISPR/CRISPRA targeting and non-targeting control OPCs were derived from the same batch of OPCs and infected and selected simultaneously. qPCR was performed to validate a reduction or overexpression of gene targets of interest for each batch of CRISPR/CRISPRA OPCs generated.

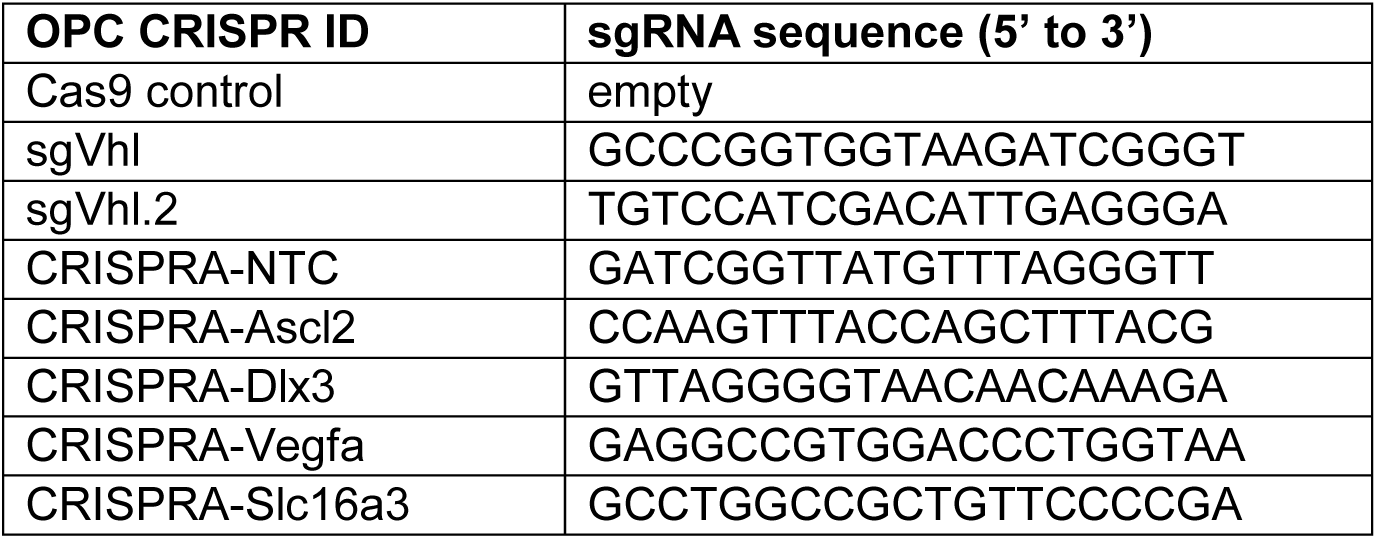

### Validating CRISPR knockout in OPCs

Primers were identified surrounding the target cut site for the two sgVhl constructs (see previous section for sgRNAs) that generate 200-250 base pair amplicons.

For the sgVhl cut site the primers were:

F 5’ TCCCTACACGACGCTCTTCCGATCTCTCTCAGGTCATCTTCTGCAACC 3’

R 5’ AGTTCAGACGTGTGCTCTTCCGATCTGACAAGATGCTCGGGGTCGG 3’

For the sgVhl.2 cut site the primers were:

F 5’ TCCCTACACGACGCTCTTCCGATCTAATAAACAGGTGCCATGCCC 3’

R 5’ AGTTCAGACGTGTGCTCTTCCGATCTAGATTGACTATTAACCTGGCAATG 3’

PCR products were run on an agarose gel, excised by gel extraction (28115, Qiagen), and submitted to the Case Western Reserve University Genomics core for library preparation and sequencing. Libraries were prepared by adding unique indices by PCR using KAPA HiFi HotStart ReadyMix. Samples were then pooled evenly, quantified using NEBNext® Library Quant Kit for Illumina® (E7630, New England 641 Biolabs), and denatured and diluted per Illumina’s MiSeq instructions. These finished libraries were then sequenced using an Illumina MiSeq (250bp paired-end). Results were analyzed using Outknocker software (Schmid-Burgk et al., 2014) (http://www.outknocker.org/outknocker2.htm) to calculate the percentage of reads with insertions or deletions at the sgRNA target site.

### Compound screening and assessment

Compound screening was carried out as described in (Lager et al., 2018). Poly-D-lysine-coated 384-well CellCarrier Ultra plates (PerkinElmer) were coated with laminin diluted in N2B27 base media using an EL406 Microplate Washer Dispenser (BioTek) equipped with a 5µl dispense cassette (BioTek) and were incubated at 37°C for at least 1 hour. A 3mM stock of the Selleck bioactive library in dimethylsulfoxide (DMSO) was then added to the plates using a 50nL solid pin tool attached to a Janus automated workstation (Perkin Elmer) at a 1:1000 dilution such that each well received a single compound at a final concentration of 3µM. Compounds for dose response validation were sourced from the Selleck library, except for ERK1/2 inhibitors SCH772984 (Selleck, S7101), AZD0364 (Selleck, S8708), and VX-11e (Selleck, S7709) which were purchased separately. OPCs were dispensed in oligodendrocyte differentiation media at 12,500 cells per well into the laminin-coated 384 well plates using the BioTek EL406 Microplate Washer Dispenser and differentiated at 37°C for 3 days. At this point, cells were fixed, washed and stained using the BioTek EL406 Microplate Washer Dispenser. Cells were stained with anti-MBP (1:100, Abcam, ab7349) along with DAPI (1 μg/ml, Sigma) and imaged using the Operetta High Content Imaging and Analysis system (PerkinElmer).

### Western blot

For cell culture derived protein samples, at least 1 million OPCs were collected and lysed in RIPA buffer (Sigma) supplemented with protease and phosphatase inhibitor (78441, Thermo Fisher) for at least 15 minutes and cleared by centrifugation at 13,000g at 4°C for 15 minutes. Protein concentrations were determined using the Bradford assay (Bio-Rad Laboratories). Protein was then diluted and added to Laemmli loading buffer, boiled at 95°C for 5 minutes, run using NuPAGE Bis-Tris gels (NP0335BOX, Thermo Fisher), and then transferred to PVDF membranes (LC2002, Thermo Fisher). Blocking and primary/secondary antibody solutions were performed for at least 30 min with 5% nonfat drug milk (Nestle Carnation) in TBS plus 0.1% Tween 20 (TBST). Primary antibodies used included anti-HIF1a (1:500, Abcam, ab2185), anti-phospho-p44/42 (ERK1/2) (100ng/mL, CST, 9101), anti-p44/42 (total ERK1/2) (100ng/mL, CST, 4696), anti-DLX3 (2µg/mL, abcam, ab178428), anti-SOX10 (1:100, R&D, AF2864), anti-B-Actin peroxidase (1:50,000, Sigma, A3854), anti-VHL (5µg/mL, BD Biosciences, 556347), anti-ASCL2 (1:10, EMD Millipore, MAB4417), anti-MBP (1µg/mL, Biolegend, 808401), and anti-MAG (1µg/mL, Thermo Fisher, 346200). Membranes were then imaged using the LI-COR (Odyssey) and analyzed using Image Studio™software that is integrated into the LI-COR imaging suite. Westerns were normalized to loading control Beta-Actin unless otherwise noted.

### qRT-PCR

At least 500,000 OPCs were lysed in TRIzol (Ambion) followed by phenol-chloroform extraction and processing with the RNeasy Mini Kit (74104, Qiagen). RNA quality and quantity was determined using a NanoDrop spectrophotometer. cDNA was generated using the iSCRIPT kit following the manufacturer’s instructions (1708891, Biorad). qRT-PCR was performed using pre-designed TaqMan gene expression assays (Thermo Fisher). Probe cat numbers and IDs are included in the chart below. qPCR was performed using the Applied Biosystems 7300 real-time PCR system and probes were normalized to *Rpl13a* endogenous control.

| Gene ID | Taqman cat. number |
| --- | --- |
| *Sox10* | Mm00569909_m1 |
| *Rpl13a* | Mm05910660_g1 |
| *Vhl* | Mm00494137_m1 |
| *Ascl2* | Mm01268891_g1 |
| *Tpi1* | Mm00833691_g1 |
| *Ciart* | Mm01255905_g1 |
| *P4ha2* | Mm00477940_m1 |
| *Pdk1* | Mm00554300_m1 |
| *Pfkl* | Mm00435587_m1 |
| *Slc16a3* | Mm00446102_m1 |
| *Dlx3* | Mm00438428_m1 |
| *Vefga* | Mm00437306_m1 |
| *Bnip3* | Mm01275600_g1 |
| *Pdgfra* | Mm00440701_m1 |
| *Plp1* | Mm01297210_m1 |
| *Myrf* | Mm01194959_m1 |

### RNA-seq sample preparation and analysis

At least 1 million OPCs were lysed in TRIzol and RNA was isolated as described for qPCR. Libraries were prepared following protocols from NEBNext Poly(A) mRNA Magnetic Isolation Module (NEB #E7490L) and NEBNext Ultra RNA Library Prep Kit for Illumina (NEB# E7530L). In brief, samples were enriched for mRNA using oligo(dT) beads, which were then fragmented randomly and used for cDNA generation and subsequent second-strand synthesis using a custom second-strand synthesis buffer (Illumina), dNTPs, RNase H and DNA polymerase I. cDNA libraries then went through terminal repair, A-base ligation, adapter ligation, size selection and PCR enrichment. Final libraries were pooled evenly and sequenced on the Illumina NovaSeq with paired-end 150bp reads with a read-depth of at least 20 million reads per sample.

For gene expression analysis, reads were aligned to the mm10 genome build using salmon 0.14.1 (https://github.com/COMBINE-lab/salmon) to quantify transcript abundance in transcripts per million (TPM) values. Transcripts were summarized as gene-level TPM abundances with tximport. A gene with TPM>1 was considered expressed. Differential expression analysis was then performed using DESEQ2 (https://bioconductor.org/packages/release/bioc/html/DESeq2.html). Significant genes were called based on p-adj and fold change values as described in the results section.

### Gene ontology analysis and heatmaps

Metascape (http://metascape.org/) was used to identify significant pathways from RNAseq data. Identification of critical oligodendrocyte genes that were dysregulated in VHL knockout OPCs (see Figure 3A) was performed by fitting RNA-seq (TPM) to the list of genes under the GO term “Oligodendrocyte Development” (GO: 0014003). This spreadsheet of TPM values was used to make a heatmap using the R-package “pheatmap.” Increased genes were those greater than 1.25 fold change whereas decreased genes were those less than 0.75 fold change in sgVhl relative to Cas9 control OPCs. In Figure 5D and 5E, RNA-seq data was fit to the genes within “Oligodendrocyte Development” (GO: 0014003) that were downregulated in sgVhl OPCs compared to Cas9 control OPCs (fold change less than 0.75).

### Gene set enrichment analysis

Gene set enrichment analysis (GSEA) scores were generated for gene sets in C5.bp datasets using classic scoring, 1000 gene-set permutations, and signal-to-noise metrics. Normalized enrichment scores, false discovery rate and FWER o-values were all calculated by GSEA software (https://www.gsea-msigdb.org/gsea/index.jsp).

### ChIP-seq and analysis

Nuclei isolation and chromatin shearing were performed using the Covaris TruChIP protocol following manufacturer’s instructions for the “high-cell” format. In brief, 5-20 million cells (H3K27Ac) or 100 million cells (HIF1a) were crosslinked in “Fixing buffer A” supplemented with 1% fresh formaldehyde for 10 minutes at room temperature with oscillation and quenched for 5 minutes with “Quench buffer E.” These cells were then washed with PBS and either snap frozen and stored at −80°C, or immediately used for nuclei extraction and shearing per the manufacturer protocol. The samples were sonicated with the Covaris S2 using the following settings: 5% Duty factor 4 intensity for four 60-second cycles. Sheared chromatin was cleared and incubated overnight at 4 degrees with primary antibodies that were pre-incubated with protein G magnetic DynaBeads (Thermo Fisher). Primary antibodies used included anti-H3K27Ac (9ug/sample, Abcam, ab4729) and anti-HIF1a (25ug/sample, Abcam, ab2185). These beads were then washed, eluted, reverse cross-linked and treated with RNase A followed by proteinase K. ChIP DNA was purified using Ampure XP beads (Aline Biosciences, C-1003-5) and then used to prepare Illumina sequencing libraries as described previously (Schmidt et al., 2009). Libraries were sequenced on the Illumina HiSeq2500 with single-end 50bp reads with a read-depth of at least 20 million reads per sample.

For peak calling, reads were quality and adapter trimmed using Trim Galore! Version 0.4.1. Trimmed reads were aligned to mm10 with Bowtie2 version 2.3.2. Duplicate reads (potential artifacts of PCR in library preparation) were removed using Picard MarkDuplicates. Peaks were called with MACS version 2.1.1 to define broad peaks for histone marks (H3K27Ac) and narrow peaks for transcription factors (HIF1a) and normalized to background genomic DNA with matched inputs. Thresholding was set at FDR<0.001 for calling both H3K27Ac and HIF1a peaks. Peaks were visualized with the Integrative Genomics Viewer (IGV, Broad Institute). Peaks were assigned to the nearest expressed gene (TPM>1 in Cas9 control or sgVhl OPCs) using bedtools available in Galaxy (https://usegalaxy.org).

### Diffbind analysis

H3K27Ac ChIP-seq was performed in duplicate from two independent batches of Cas9 control and sgVhl OPCs. Differential H3K27Ac analysis between sgVhl and Cas9 control OPCs was performed using “Diffbind software” (https://bioconductor.org/packages/release/bioc/html/DiffBind.html). A false discovery rate of 0.1 was used to call significantly enriched and depleted regions of H3K27Ac.

### Motif enrichment analysis

Motifs were called under significant HIF1a peaks (FDR<0.001) or regions of significantly gained H3K27Ac (FDR<0.1) using HOMERv4.11.1 (Heinz et al., 2010). The FindMotifsGenome.pl tool was used with 1000bp windows with mm10 as the reference genome.

### HIF1a ChIP-seq overlap analysis

HIF1a ChIP-seq raw data were re-analyzed for: two HIF1a replicates ChIP-seqs and 1 input of E12.5 heart (GSM1500750, GSM1500751, GSM1500749) (Guimaraes-Camboa et al., 2015), two HIF1a replicates ChIP-seqs and 1 input of melanocyte (GSM2305570, GSM2305571, GSM2305572) (Loftus et al., 2017), and two HIF1a ChIP-seqs and 2 inputs of Th17 cells (GSM1004819, GSM1004991, GSM1004820, GSM1004993) (Ciofani et al., 2012). For each of the HIF1a ChIP-seqs, the peaks were called as described above (MACS2, narrow peak, FDR<0.001). Peaks unique to OPCs were identified by finding peaks not found in the union of all the replicates of every other tissue sample using the bedtools (v 2.25.0) (Quinlan and Hall, 2010) “intersect” command. Common peaks across tissues were also identified using the bedtools “intersect”. RPCG-normed bigwigs were used to create aggregate plots for HIF1a, ATAC-seq, and H3K27ac using the deeptools (version 3.3.1) (Ramirez et al., 2016) “computeMatrix” and “plotHeatmap” commands, centering on the HIF1a peak locations in the sgVhl OPCs. Genes within 5Kb of these HIF1a peaks were called as HIF1a target genes. For melanocytes and T-cells, HIF1a target genes were called as only those at the intersection of both of the available replicates. For embryonic heart there were fewer peaks called compared to the other tissues, so HIF1a gene targets were called as the union of the two available replicates. These gene and peak lists were used to generate venn diagrams using the following webtool: (http://bioinformatics.psb.ugent.be/webtools/Venn/). Cell type specific gene targets of HIF1a were called if the gene was only a target of HIF1a in that specific cell type.

### Mass spectrometry for lactate and pyruvate

OPCs were cultured in OPC growth media and after growth media was removed, cells were washed with ice-cold saline (3 times) and collected with 1 ml of 80% ethanol pre-chilled on dry ice. Cells were frozen at −80°C until analyses. This cell extract was then vortexed and sonicated for 30 seconds on, 30 seconds off, alternating for 10 minutes. Next, cells were pelleted by centrifugation at 4°C for 10 min at 14,000 rpm. Supernatant was transferred to GC/MS vials and evaporated to dryness under gentle stream of nitrogen. Keto- and aldehyde groups were reduced by addition of 10 µl of 1 N NaOH plus 15 µl NaB^2^H_4_ (prepared as 10 mg/ml in 50 mM NaOH). After mixing, samples were incubated at room temperature for 1 hour and then acidified by 55 µl of 1 N HCl by dropping the acid slowly. Next, samples were evaporated to dryness. Next, 50 µl of methanol were added to precipitate boric acid. Internal standard was added (10 µl of 17:0 FA, 0.1 mg/ml). Samples were evaporated to dryness and reacted with 40 µl of pyridine and 60 µl of tert-butylbis(dimethylsilyl) trifluoroacetamide with 10% trimethylchlorosilane (Regisil) TBDMS at 60°C for 1 hour. Resulting TBDMS derivatives were injected into GC/MS. Analyses were then carried out on an Agilent 5973 mass spectrometer equipped with 6890 Gas Chromatograph. A HP-5MS capillary column (60 m × 0.25 mm × 0.25 μm) was used in all assays with a helium flow of 1 ml/min. Samples were analyzed in Selected Ion Monitoring (SIM) mode using electron impact ionization (EI). Ion dwell time was set to 10 msec. Lactate and pyruvate were both measured for all samples.

### Tissue from mild chronic hypoxia (MCH) mice

MCH is a well described model of Diffuse White Matter Injury (DWMI) (Clayton et al., 2017a; Clayton et al., 2017b; Fancy et al., 2011; Scafidi et al., 2014; Yuen et al., 2014) and protein samples were generously provided by Brian Popko. In brief, postnatal day 3 (P3) C57Bl6 pups were placed into a BioSpherix chamber maintained at 10+/- 0.5% O2 by displacement of nitrogen until P11. Animals were then quickly sacrificed by CO_2_ asphyxiation followed by decapitation and the frontal cortex was isolated as this region has been shown to contain subcortical white matter that is susceptible to hypoxia-induced DWMI (Clayton et al., 2017b; Jablonska et al., 2012; Sanjana et al., 2014; Yuen et al., 2014).

### Oligocortical spheroid generation and immunohistochemistry

Human embryonic stem cells (line H7, WiCell) were grown in mTesR1 media and oligocortical spheroids were generated as previously described (Madhavan et al., 2018). Spheroids were treated every other day with DMSO or 300nM AZD8330 between days 70 and 74 and harvested on day 90. 2 hours prior to harvesting, spheroids were treated with 200uM Hypoxyprobe-1 (pimonidazole, Hypoxyprobe Inc, Burlington MA). Spheroids were washed in PBS and fixed overnight in ice cold 4% Paraformaldehyde and then washed in PBS and cryoprotected in a 30% sucrose solution. Spheroids were frozen in OCT and sectioned at thickness of 15 µm. Slides were washed in PBS and incubated overnight with anti-MyRF (1:1000, gift from Michael Wegner) and anti-pimonidazole antibodies (1:250, Hypoxyprobe Inc, Burlington MA) followed by labelling with appropriate Alexa-Fluor labeled secondary antibodies (2µg/mL, Thermo Fisher). Images were captured using a Hamamatsu Nanozoomer S60 Slide scanner with NDP 2.0 software. Images spanning the edge to the central region of each oligocortical spheroid were used for analysis. MyRF+ oligodendrocytes in Hypoxyprobe+ and Hypoxyprobe-areas within each image were quantified using ImageJ software.

### Statistics and replicates

GraphPad Prism was used to perform statistical analyses unless otherwise noted. Statistical tests and replicate descriptions are detailed in each figure legend. In brief, black filled-in circles for bar graphs indicate biological replicates whereas open circles represent technical replicates. Statistics were only performed on samples with biological replicates. Data is typically graphed as mean ± standard deviation (SD) or ± standard error of the mean (SEM) as detailed in the figure legend. A p-value less than 0.05 was considered significant unless otherwise noted.

---

**Figure S1.**
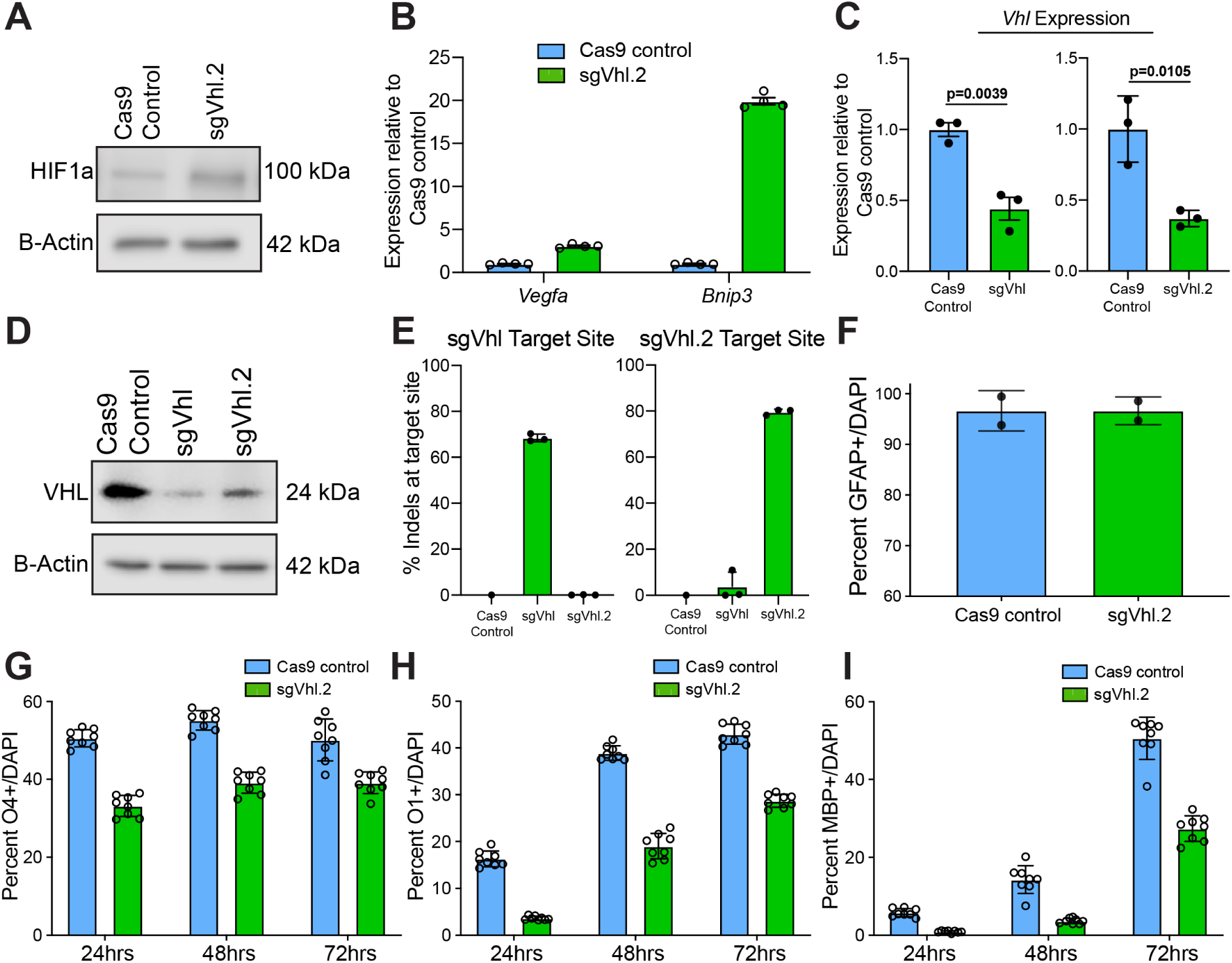
**CRISPR Mediated Knockout of Vhl Leads to Loss of *Vhl* Expression and Impairs Oligodendrocyte Formation, Related to Figure 1.** **(A)** Western blot of HIF1a from nuclear lysates of sgVhl.2 OPCs compared to Cas9 control OPCs with B-Actin as a loading control. Molecular weight is indicated to the right of the blot. **(B)** qRT-PCR of HIF target genes *Vegfa* and *Bnip3* in sgVhl.2 OPCs (in green) compared to Cas9 control OPCs (in blue) normalized to endogenous loading control *Rpl13a*. Data are presented as mean ± SEM from 4 technical replicates (individual wells). **(C)** qRT-PCR of *Vhl* in both sgVhl and sgVhl.2 OPCs (both in green) compared to Cas9 control OPCs (in blue) normalized to endogenous loading control *Rpl13a*. Data are presented as mean ± SEM from 3 independent biological replicates (experiments) with 4 technical replicates (individual wells) per experiment. p-values were calculated using Student’s two-tailed t-test. **(D)** Western blot of VHL from whole cell lysate of sgVhl and sgVhl.2 OPCs compared to Cas9 control OPCs with B-Actin as a loading control. Molecular weight is indicated to the right of the blot. **(E)** In-del analysis utilizing Outknocker software on PCR products surrounding sgRNA cut sites for both sgVhl and sgVhl.2 OPCs compared to Cas9 control. Data are presented as mean ± SD from 3 independent biological replicates (experiments). **(F)** Quantification of the percentage of astrocytes (GFAP+ cells/ DAPI) formed from sgVhl.2 OPCs (in green) compared to Cas9 control OPCs (in blue). Data are presented as mean ± SD from 2 independent biological replicates (experiments) with 6-8 technical replicates (individual wells) per experiment. **(G-I)** Quantification of the percentage of early O4+ (G), intermediate O1+ (H), and late MBP+ (I) oligodendrocytes formed from sgVhl.2 OPCs (in green) compared to Cas9 control OPCs (in blue) at day 1, 2 and 3 of differentiation. Data are presented as mean ± SD from 8 technical replicates (individual wells) per condition.

**Figure S2.**
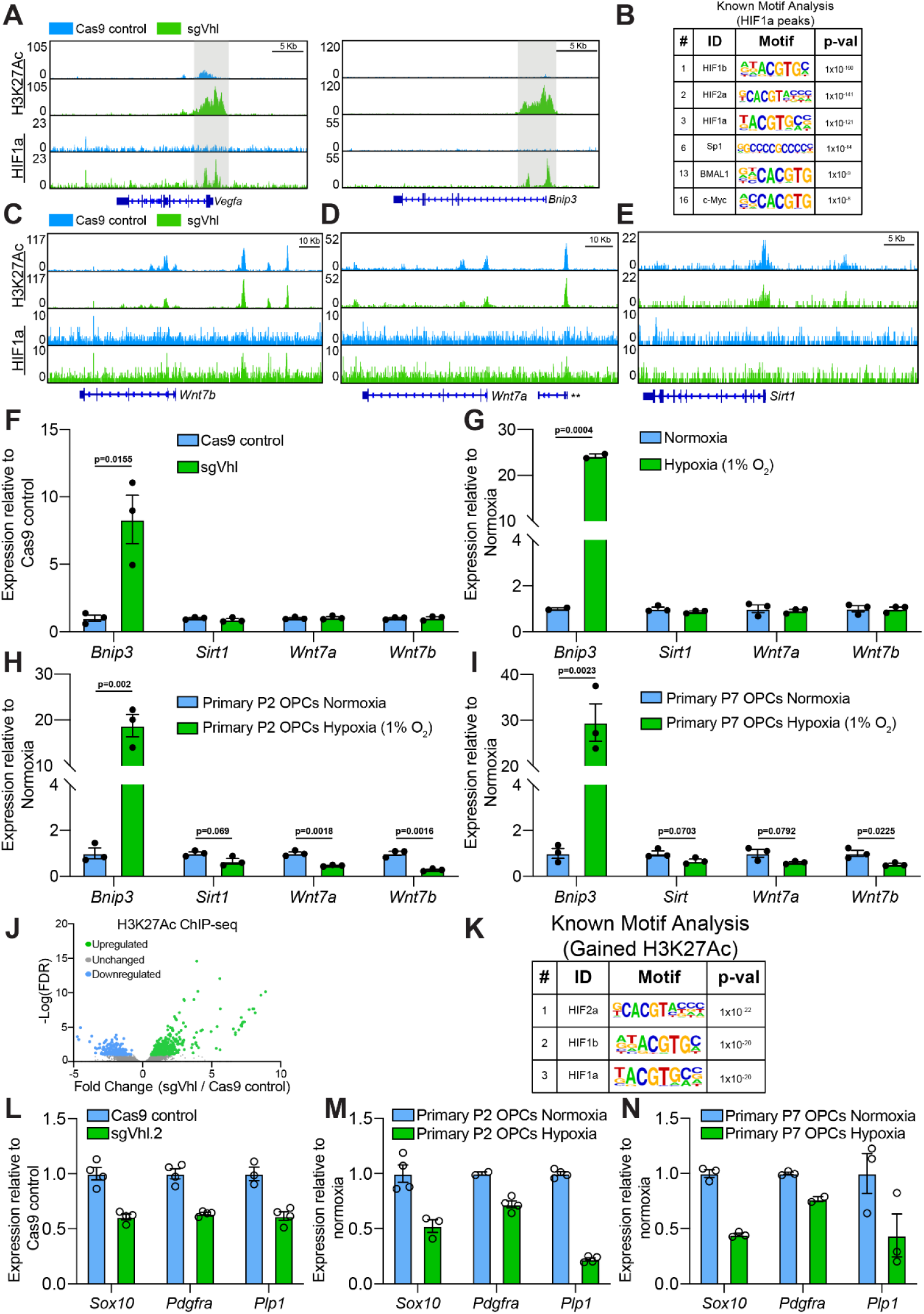
**HIF1a Accumulation Suppresses Sox10 Expression, Related to Figure 2.** **(A)** Genome browser view of H3K27ac and HIF1a signals in Cas9 control (in blue) and sgVhl (in green) OPCs normalized to input at the locus for *Vegfa* and *Bnip3*. HIF1a and active chromatin accumulation in sgVhl OPCs are highlighted in gray. Scale bars, 5Kb. **(B)** Table of known motifs significantly enriched in HIF1a peaks in sgVhl OPCs (FDR<0.001) displaying the transcription factor name, motif, and p-value ranked in order of significance (# indicates rank out of all motifs from the analysis). **(C-E)** H3K27ac and HIF1a ChIP-seq in Cas9 control (in blue) and sgVhl (in green) OPCs normalized to input at the locus for *Wnt7b*. Scale bars, 10Kb (C-D) and 5Kb (E). ** is 4930471M09Rik. **(F-I)** qRT-PCR of *Bnip3, Sirt1, Wnt7a* or *Wnt7b* in Cas9 control OPCs (in blue) and sgVhl OPCs (in green) (F), OPCs treated with normoxia (in blue) or hypoxia (1% O_2_, in green) (G), primary postnatal day 2 (P2) *in vivo* derived mouse OPCs cultured in physiological hypoxia (1% O_2_, in green) compared to normoxia (in blue) (H), and primary immunopanned postnatal day 7 (P7) *in vivo* derived mouse OPCs cultured in physiological hypoxia (1% O_2_, in green) compared to normoxia (in blue) (I) normalized to endogenous loading control *Rpl13a*. Data are presented as mean ± SEM from 3 independent biological replicates (experiments) with 4 technical replicates (individual wells) per experiment. p-values were calculated using Student’s two-tailed t-test. **J)** Volcano plot of fold change in intensity of H3K27ac peaks (FDR<0.001) between Cas9 control and sgVhl OPCs showing regions of significantly increased H3K27ac (FDR<0.1, in green) and decreased H3K27ac (FDR<0.1, in blue). **(J)** Volcano plot of fold change in intensity of significant H3K27ac regions (FDR<0.001) between Cas9 control and sgVhl OPCs showing regions of significantly increased H3K27ac (FDR<0.1, in green) and decreased H3K27ac (FDR<0.1, in blue). **(K)** Table of known motifs in regions significantly enriched for H3K27ac (FDR<0.1) in sgVhl OPCs compared to Cas9 control OPCs displaying the transcription factor name, motif, and p-value ranked in order of significance (# indicates rank out of all motifs from the analysis). **(L-N)** qRT-PCR of *Sox10* and downstream Sox10 target genes *Pdgfra* and *Plp1* in sgVhl.2 (in green) compared to Cas9 control OPCs (in blue) (L), in primary postnatal day 2 (P2) *in vivo* derived mouse OPCs treated with physiological hypoxia (in green) compared to normoxia (in blue) (M), and in primary immunopanned postnatal day 7 (P7) *in vivo* derived mouse OPCs treated with physiological hypoxia (in green) compared to normoxia (in blue) (N). All qRT-PCRs are normalized to endogenous loading control *Rpl13a*. Data are presented as mean ± SEM from 3-4 technical replicates (individual wells) per condition.

**Figure S3.**
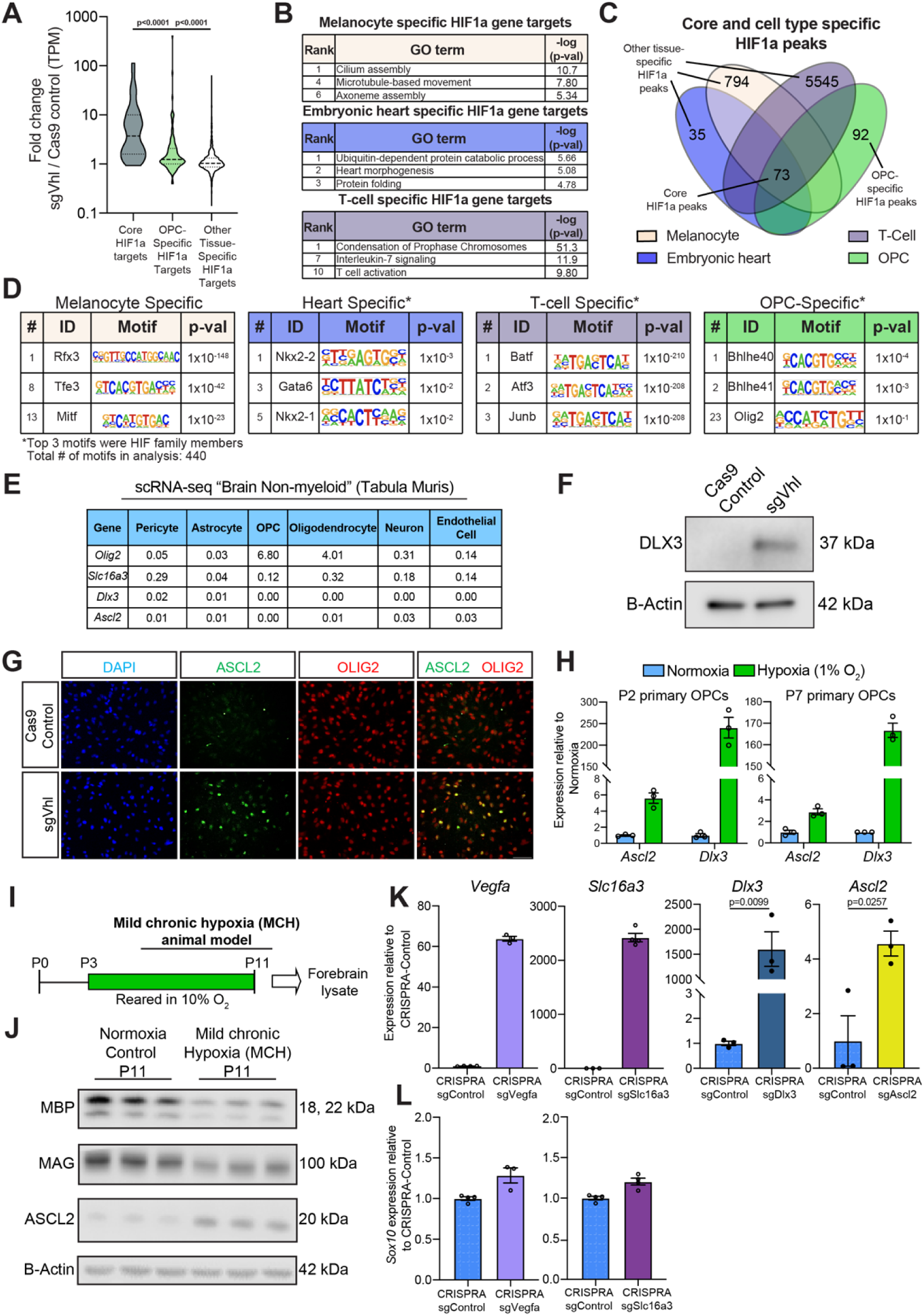
**OPC-specific HIF1a Targets Downregulate Sox10, Related to Figure 3.** **(A)** Violin plot of fold change in expression (TPM) between Cas9 control and sgVhl OPCs of genes included in core (in dark gray), OPC-specific (light green) and other tissue-specific (light gray) HIF1a target categories. Bold dashed line represents the median with the thin dashed lines representing the upper and lower quartiles. p-values were calculated using the Kruskal Wallis One-Way ANOVA with Dunn’s multiple comparisons test. **(B)** Gene ontology (GO) analysis of genes targeted by HIF1a specifically in melanocytes (beige), embryonic heart (purple), and T-cells (magenta). The chart includes curated pathways with their rank based on their respective p-values. **(C)** Overlap of significant HIF1a peaks (FDR<0.001) across 4 different cell types giving core HIF1a peaks, OPC-specific HIF1a peaks and other tissue-specific HIF1a peaks (combination of heart, T-cell and melanocyte specific peaks). **(D)** Table of known motifs significantly enriched in cell-type-specific HIF1a peaks in melanocytes (beige), heart (purple), T-cells (magenta) and OPCs (light green). Charts display the transcription factor name, motif, and p-value ranked in order of significance (# indicates rank out of all 440 motifs in the analysis and * indicates that HIF motifs were removed). **(E)** Chart of publicly available single-cell RNA-seq data from non-myeloid cells of the brain for expression of *Dlx3, Ascl2* and *Slc16a3* with positive control *Olig2*, which marks OPCs and oligodendrocytes. Values are ln(1+CPM). CPM is counts per million reads. **(F)** Western blot of nuclear lysates for DLX3 in sgVhl OPCs and CRISPR-control OPCs with B-Actin as the loading control. Molecular weight is indicated to the right of the blot. **(G)** Immunocytochemistry (ICC) for ASCL2 (in green) and OLIG2 (in red) in Cas9 control and sgVhl OPCs. Nuclei are marked by DAPI (in blue). **(H)** qRT-PCR of *Dlx3* and *Ascl2* in P2 and P7 primary *in vivo* derived OPCs treated with physiological hypoxia (1% O_2_, in green) compared to normoxia (21% O_2_, in blue) normalized to endogenous loading control *Rpl13a*. Data are presented as mean ± SEM from 3 technical replicates (individual wells) per condition. **(I)** Schematic highlighting the mild chronic hypoxia model of diffuse white matter injury in which mouse pups are reared in 10% oxygen from postnatal day 3 to 11 (P3 to P11) to model diffuse white matter injury of prematurity. Animals are sacrificed at postnatal day (P11) and their forebrains are isolated and lysed for protein. **(J)** Western blots of forebrain lysates from P11 female C57Bl6 mice of myelin proteins MBP and MAG along with ASCL2 in animals reared in hypoxia from P3-P11 compared to control normoxic reared mice. B-Actin is shown as a loading control. N= 3 female C57Bl6 mice per treatment group. Molecular weight is indicated to the right of the blot. **(K)** qRT-PCR of *Vegfa, Slc16a3, Ascl2* and *Dlx3* normalized to endogenous loading control *Rpl13a* in their respective CRISPRA OPCs compared to CRISPRA sgControl OPCs. Data are presented as mean ± SEM from 3-4 technical replicates (individual wells) per condition. **(L)** qRT-PCR of *Sox10* in CRISPRA sgVegfa (light purple) and CRISPRA sgSlc16a3 (in magenta), OPCs compared to CRISPRA sgControl OPCs normalized to endogenous loading control *Rpl13a*. Data for CRISPRA sgVegfa and CRISPRA sgSlc16a3 are presented as mean ± SEM from 3-4 technical replicates (individual wells) per condition.

**Figure S4.**
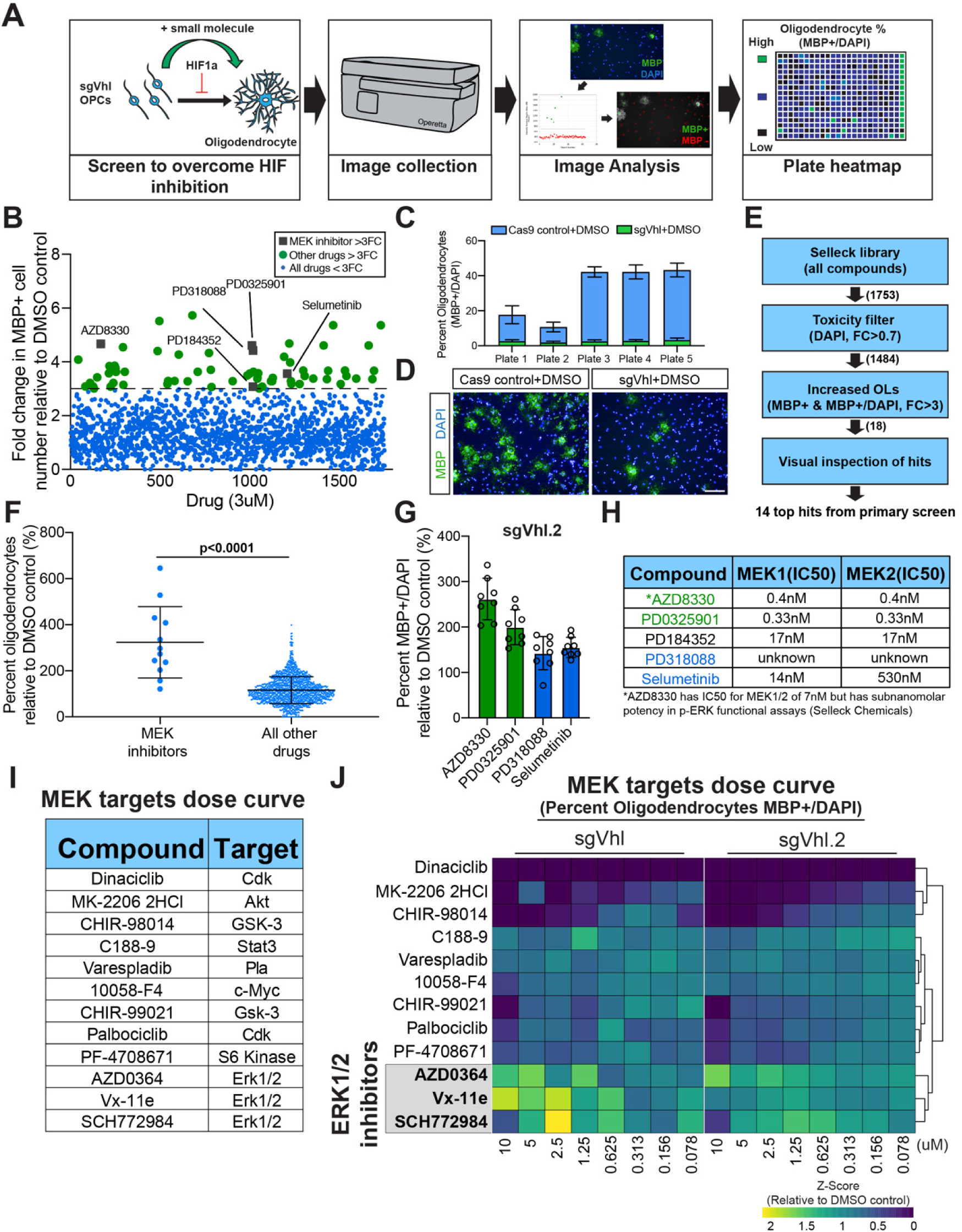
**Inhibition of MEK/ERK Signaling Increases Oligodendrocyte Formation from sgVhl OPCs, Related to Figure 4.** **(A)** Schematic depicting the procedure for the primary bioactives screen to uncover compounds that increase oligodendrocyte formation from sgVhl OPCs. **(B)** Primary bioactives library screen showing the effect of 1753 small molecules on number of oligodendrocytes (MBP+ cells) formed by sgVhl OPCs relative to DMSO treated sgVhl OPCs. Anything higher than the dotted line represents a greater than 3-fold change increase from DMSO (green dots). MEK inhibitors are highlighted as gray boxes with their respective drug names. **(C)** Primary screen positive control (Cas9 control+DMSO) and negative control (sgVhl+DMSO) percent oligodendrocyte (MBP+/DAPI) metrics on a per plate basis. Data represent mean ± SD from 16 technical replicates (individual wells) per plate per condition. **(D)** Representative immunocytochemistry images of oligodendrocytes (MBP+ in green) from primary screen positive (Cas9 control+DMSO) and negative (sgVhl+DMSO) controls. Nuclei are marked by DAPI (in blue). Scale bars, 100µm. **(E)** Schematic detailing the filtering steps starting with the bioactives library and narrowing down to top hits that are non-toxic (total DAPI FC<0.7), increased the number and percentage of oligodendrocytes (FC>3) and passed visual inspection to give the top 14 compound hits. Numbers in parentheses represent the number of drugs after each filtering step. **(F)** Quantification of the effect of all non-toxic MEK inhibitors (n=12) compared to all other non-toxic drugs from the primary screen (n=1472) on the percentage of oligodendrocytes from sgVhl OPCs relative to DMSO treated sgVhl OPCs. Data are presented as mean ± SD. p-values were calculated using the Mann-Whitney t-test. **(G)** Collapsing all tested doses into one overall average shows the ability of each MEK inhibitor to increase the formation of oligodendrocytes (MBP+/DAPI) from sgVhl.2 OPCs relative to DMSO treatment. AZD8330 and PD0325901 are shown in green representing the most effective drugs, while PD318088 and Selumetinib are shown in blue as slightly less effective drugs. Data are presented as the mean ± SD of all 8 doses for each drug from a single dose curve plate. **(H)** Table of IC50 values for MEK1 and MEK2 for the top MEK inhibitors AZD8330, PD035901, PD184352, and Selumetinib. IC50 is still currently unknown for PD318088. **(I)** List of the 12 drugs included on the MEK targets dose curve plate including the drug name and canonical target of the drug. **(J)** Heatmap representation of the 8-point dose curve performed for all 12 drugs in the MEK target dose curve plate shown as row Z-score. Data in the heatmap is the fold change in percent oligodendrocytes (MBP+/total DAPI) in sgVhl and sgVhl.2 OPCs relative to their respective DMSO negative controls. Rows were sorted by unsupervised hierarchical clustering and columns are in order from high (10µM) to low dose (78nM) of drug. ERK1/2 inhibitors are highlighted in gray and bold. Data are presented as the mean for each drug at each dose from 2 separate dose curve plates for both sgVhl and sgVhl.2 OPCs.

**Figure S5.**
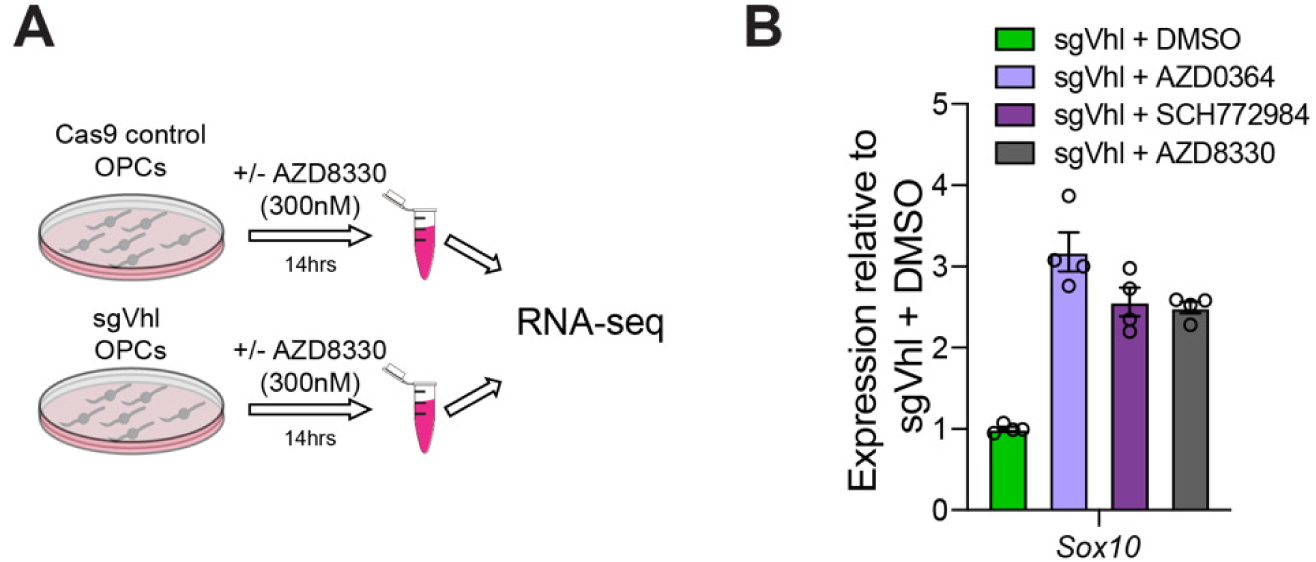
**Inhibition of MEK/ERK Signaling Drives *Sox10* Expression in sgVhl OPCs, Related to Figure 5.** **(A)** Schematic highlighting the setup of the RNA-seq experiment in which Cas9 control and sgVhl OPCs were treated with either DMSO or 300nM of AZD8330 for 14 hours and then cells were lysed for poly-adenylated mRNA extraction and sequenced. **(B)** qRT-PCR of *Sox10* in sgVhl OPCs treated with DMSO, AZD0364 (1µM, in purple), SCH772984 (1µM, in magenta) or AZD8330 (300nM, in gray) for 14hrs normalized to endogenous control *Rpl13a*. Data are presented as mean ± SEM from 4 technical replicates (individual wells) per condition.

## Notes

### Summary of Updates

Correct image added to Figure 4C

